# Behavioral strategy shapes activation of the Vip-Sst disinhibitory circuit in visual cortex

**DOI:** 10.1101/2023.04.28.538575

**Authors:** Alex Piet, Nick Ponvert, Douglas Ollerenshaw, Marina Garrett, Peter A. Groblewski, Shawn Olsen, Christof Koch, Anton Arkhipov

## Abstract

In complex environments, animals can adopt diverse strategies to find rewards. How distinct strategies differentially engage brain circuits is not well understood. Here we investigate this question, focusing on the cortical Vip-Sst disinhibitory circuit. We characterize the behavioral strategies used by mice during a visual change detection task. Using a dynamic logistic regression model we find individual mice use mixtures of a visual comparison strategy and a statistical timing strategy. Separately, mice also have periods of task engagement and disengagement. Two-photon calcium imaging shows large strategy dependent differences in neural activity in excitatory, Sst inhibitory, and Vip inhibitory cells in response to both image changes and image omissions. In contrast, task engagement has limited effects on neural population activity. We find the diversity of neural correlates of strategy can be understood parsimoniously as increased activation of the Vip-Sst disinhibitory circuit during the visual comparison strategy which facilitates task appropriate responses.

## Introduction

Circuitry across the brain, including sensory cortex, does not operate in isolation but rather serves behavioral demands. As such, quantitative behavioral analysis is an essential step in detailed understanding of brain circuits (Carandini 2012; Gomez-Marin et al. 2014; Krakauer et al. 2017; Niv 2021). Recently a suite of computational tools has emerged for precise analysis of behavioral tasks used in neuroscience laboratories including the description of behavioral strategies (Brunton et al. 2013; Berman 2018; Roy et al. 2018; Roy et al. 2021; Ashwood et al. 2022; Jha et al. 2022; Le et al. 2022). These tools vary in their statistical structure but all parameterize a space of possible behaviors which are then constrained by behavioral data. This has lead to a greater appreciation of behavioral diversity during laboratory tasks, including across subject variability, and within subject variability across individual behavioral sessions and task learning.

Behavioral strategies have been shown to alter which brain regions and pathways are active during a task. Gilad et al. 2018 found mice used either an active or passive whisking strategy which altered the location of short term memory in the cortex. Bolkan et al. 2022 found that both task difficulty as well as the subject’s behavioral state determined how striatal pathways influenced behavior. Other studies have demonstrated strategy dependent changes in neural dynamics across human fMRI (Venkatraman et al. 2009; Yang et al. 2023), rodent wide-field imaging (Pinto et al. 2019), and cellular activity in the fruit fly (Calhoun et al. 2019). Prefrontal structures, including the anterior cingulate cortex and medial prefrontal cortex, have been proposed to track strategy preferences and control strategy execution (Tervo et al. 2021; Domenech et al. 2020; Schuck et al. 2015; Proskurin et al. 2022). Behaviorally relevant signals have also been shown to modify activity in sensory cortex (Tseng et al. 2022) including visual flow (Leinweber et al. 2017; Schneider 2020), amplifying task relevant signals (Kim et al. 2020), dynamically scaling the range of stimulus encoding (Waiblinger et al. 2022), and value signals (Banerjee et al. 2020).

Simultaneously, there has been growing appreciation for the role neural cell types play in mediating specific computations (Pinto et al. 2015; Sylwestrak et al. 2022). Merging lines of evidence suggest a disinhibitory circuit between vasoactive intestinal peptide-positive (Vip) interneurons, somatostatin-positive (Sst) interneurons, and excitatory cells (Pfeffer et al. 2013; Kullander et al. 2021; Campagnola et al. 2022; Karnani et al. 2016). The circuit is disinhibitory, whereby Vip neurons disinhibit pyramidal activity through inhibition of Sst inhibitory neurons (Kamigaki 2019). In the visual cortex, the Vip-Sst disinhibition circuit has been implicated in a variety of computational mechanisms. Millman et al. 2020 found that Vip-Sst antagonism controlled the dynamic range of stimulus contrast for excitatory cells. Keller et al. 2020 found that the Vip-Sst disinhibitory circuit mediates the influence of visual context on excitatory cells. Fu et al. 2014 found that running in mice activates the Vip-Sst disinhibition of excitatory cells. Finally, anatomical (Williams et al. 2019; Kamigaki 2019; Ma et al. 2021) as well as functional studies (Pi et al. 2013) demonstrate that top-down feedback is preferentially routed through Vip neurons, influencing local circuits through Vip disinhibition.

Despite recent progress in analysis of behavior and cell type circuit dissection, it remains unclear how behavioral strategies influence cortical circuits through specific cell types. Given the range of circuit functions mentioned above, the Vip-Sst disinhibitory circuit is a promising substrate for mediating behavioral strategies. Here we investigate how the Vip-Sst circuit in visual cortex mediates strategy dependent demands. We used the recently collected Allen Institute Visual Behavior - 2p calcium imaging dataset to examine how the activity of Vip, Sst, and excitatory cells depend on strategy preferences during a change detection task (Garrett et al. 2023; brain-map.org). This dataset, collected with the Allen Brain Observatory experimental pipeline, contains two-photon calcium imaging in genetically defined cell types collected while mice performed a visual change detection task. This large systematic survey contains behavior from 376 imaging sessions from 82 mice thus permitting analysis of behavioral diversity. We used a dynamic logistic regression model (Roy et al. 2018) to identify strategies used by mice on this task. We find individual mice have stable strategy mixtures of a visual comparison strategy and a statistical timing strategy. Vip, Sst, and excitatory cells recorded from mice predominantly using each of these strategies show dramatic differences in activity. Further, the effects of behavioral strategy were independent from task engagement and stimulus novelty. Finally, we show that strategy differences can be succinctly described by the degree of activation of the Vip-Sst disinhibitory circuit.

## Results

### Visual change detection task

To examine the relationship between behavioral strategies and cortical circuits, we analyzed the diversity of behaviors in the Allen Institute Visual Behavior - 2p calcium imaging dataset (brain-map.org). This public dataset contains 2-photon calcium imaging from transgenic mice expressing the calcium indicator GCaMP6f in excitatory neurons (Slc17a7-IRES2-Cre;Camk2a- tTa;Ai93), Sst inhibitory neurons (Sst-IRES-Cre;Ai148), and Vip inhibitory neurons (Vip-IRES-Cre;Ai148). The data was collected with the Allen Brain Observatory experimental pipeline which uses standardized data collection and processing (Garrett et al. 2023; de Vries et al. 2020). We focused our analysis on recordings from two visual areas, V1 and LM, at multiple cortical depths. During imaging, mice performed a visual change detection task (Fig. 1A). In this task, head fixed mice were shown a series of natural images (250ms stimulus duration) interspersed with periods of a gray screen (500ms inter-stimulus duration). This task uses a roving baseline paradigm whereby an individual image repeats a variable number of times before a new image is presented and then itself repeats. Mice were given a water reward for licking in response to image changes. Premature licking delayed the time of the next image change. Further, to serve as distractors, 5% of image repeats were omitted and replaced with a continuation of the gray screen for the same duration as image presentations. Image omissions therefore did not disrupt the rhythmic nature of the stimulus. Image changes as well as the image immediately before the change were not omitted. Mice performed the task on a running disk and were free to run or not run. Running had no bearing on the task. These mice learned the task through a standardized training pipeline and were selected for imaging based on a consistent criteria of minimum task performance (Garrett et al. 2023; Groblewski et al. 2020).

**Figure 1:**
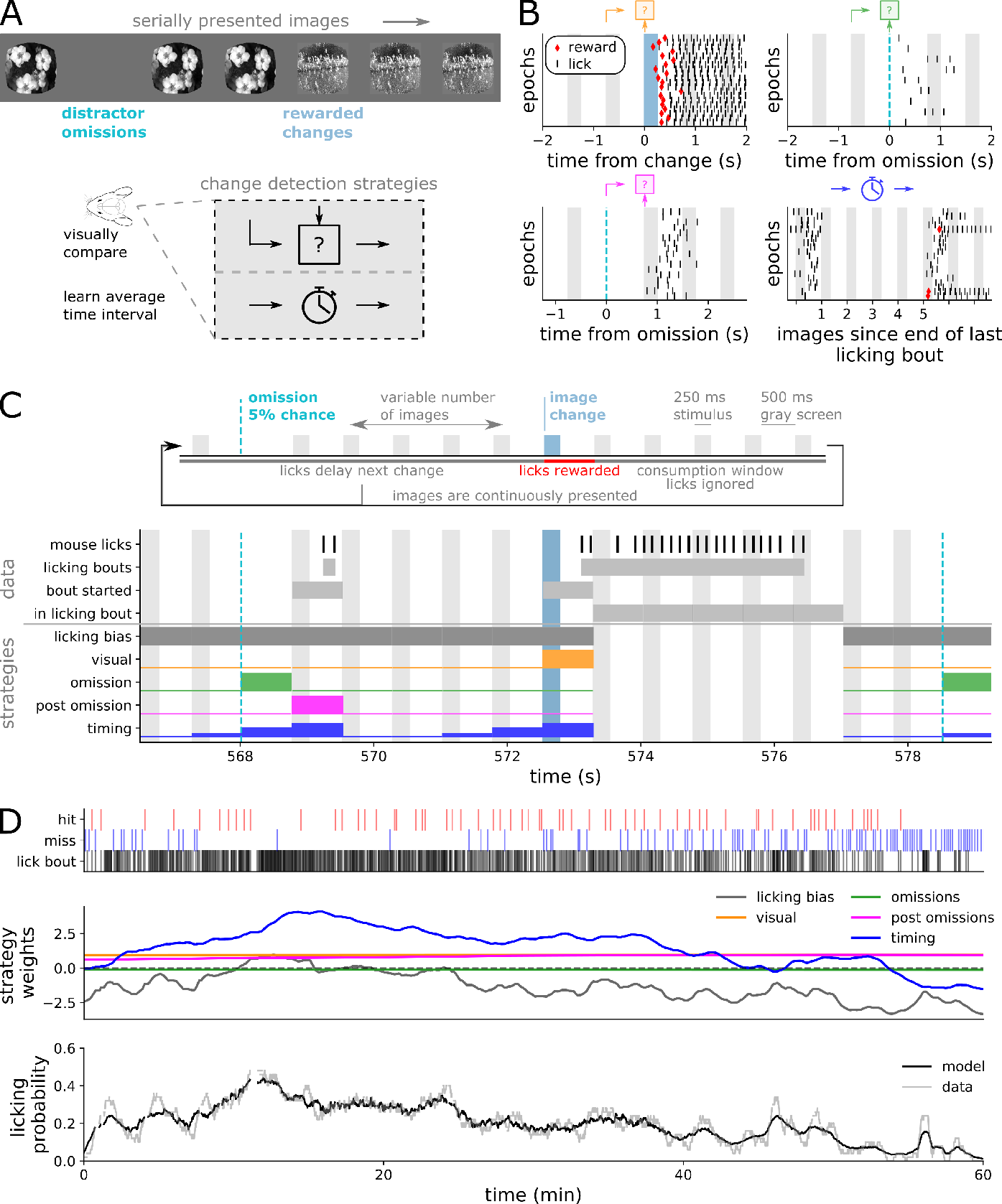
**Quantifying strategies during a change detection task**. (A) Head-fixed mice were shown a stream of natural images and were rewarded for licking in response to image changes. Mice could perform this task using multiple strategies, including visually comparing image presentations or learning statistical distributions of rewards. (B) Example lick rasters demonstrate multiple strategies. Each row is an epoch within one example session. Up to 20 examples for each strategy are shown. Gray bands show time of repeated image presentations. Blue bands show time of change image presentations. Dashed blue lines show time of omitted images. Red markers show time of rewards. Top left - licking aligned to image changes. Top right - licking aligned to image omissions. Bottom left - licking aligned to post-omission images. Bottom right - licking aligned to a fixed time interval from the last licking bout. (C) Diagram of task structure, data processing, and strategies. Images were presented for 250ms with 500ms gray screens interleaved. 5% of all images were randomly omitted. Image changes were drawn from a geometric distribution. Individual licks were segmented into licking bouts. Licking bouts were assigned to the preceding image presentation. The licking model predicts whether a licking bout starts during each image interval, and we therefore ignore image presentations where the mouse was already licking. For each strategy we show the probability of starting a licking bout during each image interval. (D) Top - Raster of licking bouts, hits, and misses for full 1-hour behavioral session. Middle - time-varying strategy weights for each strategy for this example session. Bottom - licking probability in the data and model prediction smoothed with a one minute boxcar.

Mice could potentially perform this task in multiple ways. In one possible strategy, mice could hold a memory of the previous image and perform a visual comparison to the subsequent image. This visual comparison strategy may be vulnerable to distraction from image omissions. Alternatively, mice could learn the statistical distribution of when images change and make educated guesses based on the time since the last image change (Fig. 1A). To illustrate these strategies we can examine licking patterns around time points relevant to each strategy (Fig. 1B). For one example session we can see: (top left) licking aligned to image changes, which results in a water reward, (top right) licking during image omissions, which never results in a reward, (bottom left) licking on the image after an omission, which never results in a reward, and (bottom right) licking on the fifth image after the last lick, which sometimes results in a water reward. Supplemental Note 1 shows distributions of mouse behavior across the dataset.

### Behavioral strategy model

In order to identify these strategy patterns across our dataset we used a dynamic logistic regression model (Roy et al. 2018). Our model predicts whether the mouse will start a licking bout in response to each image based on the weighted influence of each strategy. Importantly, the strategy weights were allowed to vary across the course of the session constrained by a smoothing prior described in detail below.

We will now describe how we processed the data and constructed our strategy model (Fig. 1C). First, individual mouse licks were segmented into licking bouts based on an inter-lick interval of 700ms (Supplemental Note 2A). The duration of licking bouts was largely governed by whether the mouse received and then consumed a water reward (Supplemental Note 2B). Therefore, we focused our analysis on the start of each licking bout. Licking bout onsets were time-locked to image presentations, thus for each licking bout we identified the last image or omission presented before the bout started (Supplemental Note 2C). Since our model predicts the start of licking bouts, we ignore image presentations when the mouse was already engaged in a licking bout. The design matrix of our strategy model is composed of vectors that describe the probability each strategy would start a licking bout on each image presentation. For the licking bias strategy, which is simply a bias term, the licking probability is 1 on all images representing a constant drive to lick. For the visual strategy, the probability is 1 on image changes, and 0 otherwise. For the omission strategy, the probability is 1 during image omissions, and 0 otherwise. The post-omission strategy has a probability of 1 during the first image after an omission, and 0 otherwise. For the timing strategy, the licking probability is a sigmoidal function of how many images have been presented since the end of the last licking bout. The licking probability is low immediately after a licking bout, rises to 0.5 at 4 images after a licking bout, and then a high licking probability at longer durations. The parameters of the sigmoidal function were learned from a subset of the data (Supplemental Note 3). All strategies other than the licking bias were mean-centered.

Using standard, non-dynamic, logistic regression our strategy model would predict the probability a mouse licked on a given image presentation, p(x⃗_i_), by using fixed weights, ^β⃗^, to combine the strategy vector for that image, x⃗_i_, and passing that sum through a logistic function:

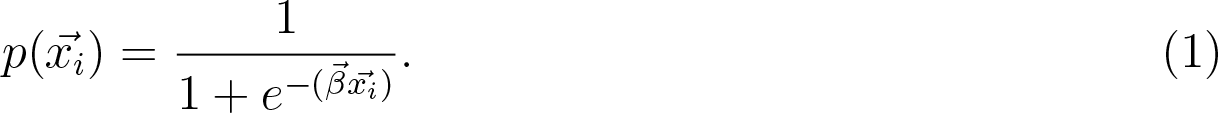

However, by using the dynamic logistic regression model, developed in Roy et al. 2018, our model lets the weight for each strategy, k, vary for each image presentation subject to a smoothing prior. The prior is implemented as letting the weights for each strategy undergo a random walk with standard deviation σ_k_:

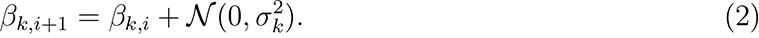

The smoothing prior σ_k_ is a hyper-parameter unique to each strategy during each session and controls the volatility of each strategy. These hyper-parameters were fit to the behavioral data by maximizing the model evidence as described in Roy et al. 2018. The smoothing prior constrains the strategy weights to evolve gradually over time. This balances the ability of the model to dynamically track changing behavioral patterns against over-fitting to the responses on each stimulus. The dynamic model has the same form as the standard model with the weights now a function of each image, ^β⃗^_i_:

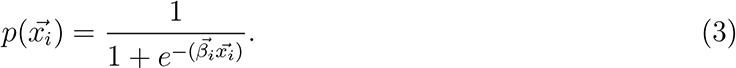

The strategy model was fit to each one-hour behavioral session by the maximum a posterori (MAP) estimate of the weights given the data and the hyper-parameters. Figure 1D shows the example output of the model for one session. For this session the licking bias and timing strategies have more volatile weights than the visual, omission, and post-omission strategies. The model successfully captures the time-varying probability of licking in the data as the mouse goes through epochs of hits, misses, as well as high and low licking rates.

### Mouse behavior is largely described by the visual and timing strategies

We quantified the performance of the model with the area under the receiver operating characteristic (ROC) curve (Fig. 2A). In this analysis we use the model’s cross validated licking probability prediction on each image as a classifier of whether the mouse started a licking bout on each image presentation. The ROC curve computes the true positive rate and false positive rate of a classifier as the classification threshold is varied. The area under this curve (AUC) ranges from .5 (classifier at chance), to 1 (perfect classification across all thresholds). The static, or standard, logistic regression model performs poorly, often at chance. The dynamic model performs well, with an average AUC value of 0.83. See Supplemental Note 4 for additional model validation details.

**Figure 2:**
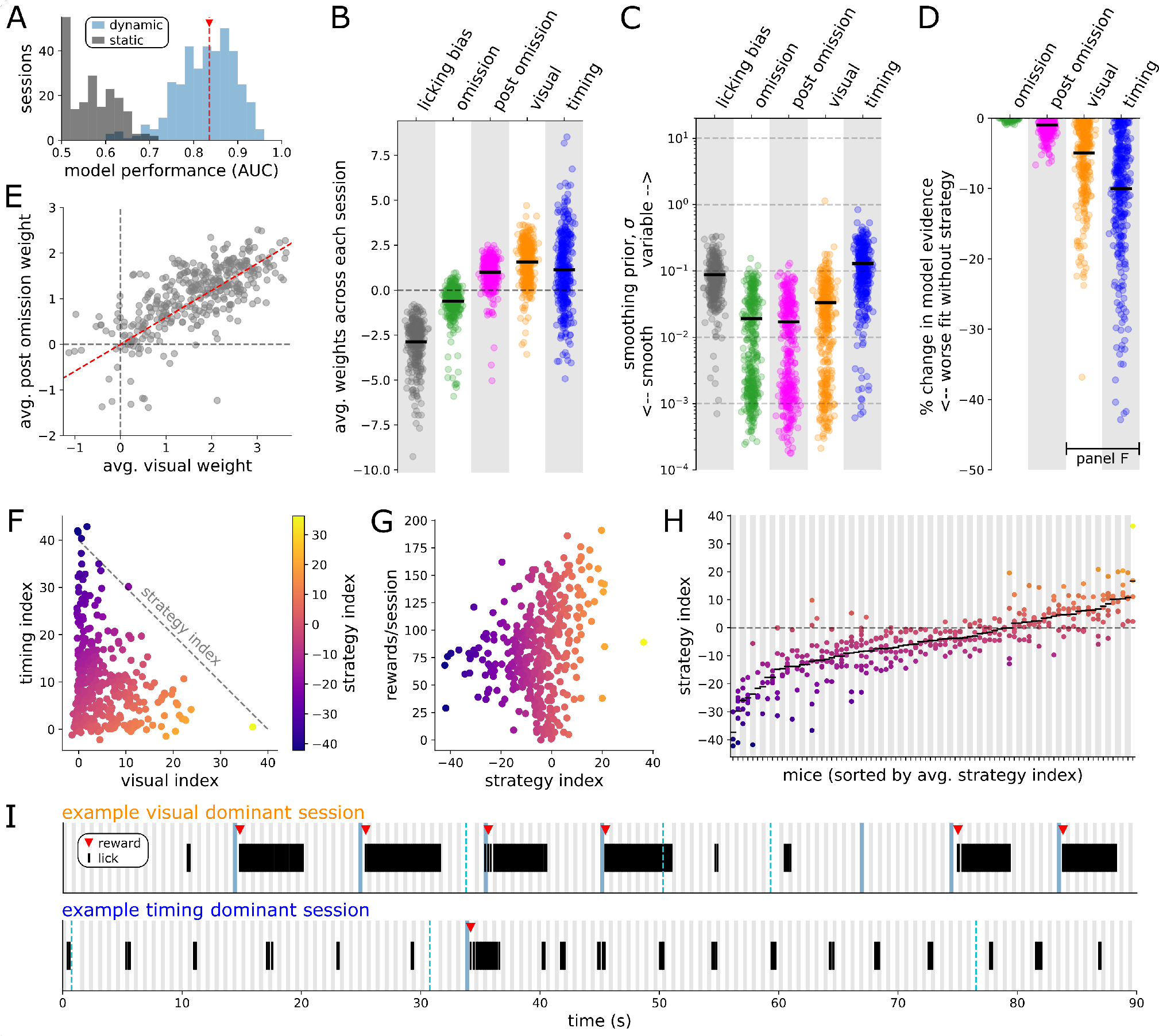
Licking model reveals distinct task strategies. (A) Cross validated model performance. Histogram of area under the ROC curve for the dynamic model (blue) and static model (gray) for each session (n=382). The red line marks the average dynamic model performance (0.83). (B-H) Dots indicate individual sessions (n=382), and (B-D) black bars are population averages. (B) Average strategy weights. (C) Learned smoothing prior σ for each strategy. (D) Reduction in model evidence when removing each strategy. The absolute value for the visual and timing strategies is shown in panel F. (E) Average weights of the visual and post-omission strategies. Red line shows a linear correlation (R^2^ = 0.44). (F) Scatter plot of the absolute value of the reduction in model evidence (termed here as an index) for the visual and timing strategies. The strategy index is defined as the counter-diagonal difference between visual and timing indices. (G) Rewards per session compared with the strategy index. (H) Mice were sorted by their average strategy index. Each session from a mouse is shown in the same column. (I) 90 seconds of illustrative behavior for two example sessions with either a visual dominant strategy (top) or timing dominant strategy (bottom). Gray bands show image repeats, blue bands mark image changes, and dashed blue lines mark image omissions.

For each behavioral session we can analyze the best fitting strategy weights (Fig. 2B) and the smoothing prior for each strategy (Fig. 2C). The weights are multiplied against the strategy design matrix and then passed through the logistic function. The licking bias strategy sets the average licking probability on each image presentation. A licking bias weight of 0 would translate to a 50% licking probability, with smaller weights leading to lower licking probabilities, and larger weights leading to higher licking probabilities. The other strategies are mean-centered (the sum of their strategy design vectors equals 0), which means we can interpret the strategy weights relative to the licking bias term. A weight of 0 would have no influence on the licking probability, a negative weight would result in less licking than the bias term, and a positive weight would result in more licking than the bias term. Across our population, the omission strategy has a negative value, meaning on average mice are less likely to lick during image omissions. The other three strategies have, on average, positive values, meaning mice are more likely to lick on the image after an omission (post omission strategy), on image changes (visual strategy), and at the expected image change frequency (timing strategy). However there is significant variability across the behavioral sessions. Supplemental Note 5 shows additional characterization of how strategy weights are correlated with the number of hits, misses, and licking probability within each session. The smoothing priors for each strategy govern how variable the strategy weights can be within each session from image to image, and constrain the strategy weights to evolve gradually over time. On average the strategies have smoothing priors within the same order of magnitude, with the licking bias and timing strategies generally being the most variable. However, there is significant variability across behavioral sessions.

To evaluate the importance of each strategy, we measured the reduction in model evidence after removing each strategy (Fig. 2D). The model evidence, or marginal likelihood, measures the probability of the data given the hyper-parameters after integrating over possible parameter values. Model comparison metrics such as Bayes factor and Bayesian information criterion are based on comparing the model evidence of two models. In general, if the model evidence decreases after removing a strategy then the model performs worse at describing the data. We did not evaluate the model evidence without the licking bias term because it sets the average licking rate and removing it breaks the model in a trivial manner. Across our population, removing the omission strategy does not lead to a reduction in model evidence. We therefore conclude that the omission strategy is not a meaningful descriptor of mouse behavior. Removing the post omission strategy leads to a small decrease in model evidence, demonstrating a minor role in describing mouse behavior. Removing the visual and timing strategies leads to significant decreases in model performance, albeit with large variability across behavioral sessions.

We focused the rest of our analysis on the visual and timing strategies based on three observations. First, the lack of change in model evidence for the omission strategy. Second, we observed a strong correlation between the post omission strategy and visual strategy in terms of both the changes in model evidence, and their average weights (Fig. 2E). Third, after performing PCA on the matrix of changes in model evidence, we found the top two principal components explained 99.04% of the variance and are closely oriented with the timing and visual strategies respectively (Supplemental Note 6).

Plotting the change in model evidence from the visual and timing strategies against each other we find that there is a continuous spectrum of behaviors that mix the visual and timing strategies together (Fig. 2F). We term the strategy index as the difference in the change of model evidence between the visual and timing strategies. A positive strategy index value indicates the session was well described by the visual strategy. A negative strategy index indicates the session was well described by the timing strategy. Plotting the strategy index against the rewards earned per session we see that all strategy mixes are able to earn a significant number of rewards per session (Fig. 2G). However higher values of the strategy index tend to result in higher number of earned rewards. Individual mice were fairly stable in their strategy preferences across multiple behavioral sessions (Fig. 2H, up to 4 sessions per mouse performed during calcium imaging). Mouse identity explains 72% of the variance in the strategy index across imaging sessions. Consistent with this finding we did not observe mice switching between strategies within a behavioral session. Further, strategy preferences emerged gradually over training (Supplemental Note 7). Taken together we find individual mice develop unique strategy preferences between the visual and timing strategies that are stable over many days.

Given that strategy preferences are stable over multiple days, the remaining analyses categorize each behavioral session by the dominant strategy (equivalently, the sign of the strategy index). We refer to sessions best described by the visual strategy (positive strategy index) as visual strategy sessions. Likewise, we refer to sessions best described by the timing strategy (negative strategy index) as timing strategy sessions. Figure 2I illustrates 90 seconds of behavior from each of the two dominant strategies. Supplemental Note 8 provides additional characterization of the behavior for each of the two strategies.

### Task strategy is distinct from task engagement

Rather than switching strategies during a session, we observed that mice had clear patterns of disengagement when they stopped licking altogether. To demonstrate this we generated a contour plot of the licking bout rate and reward rate aggregated across all the behavioral sessions (Fig. 3A). Mouse behavior is clearly divided between two regions. One region, which we term disengaged behavior, has low licking rates and low reward rates. The other region, which we term engaged behavior, is much broader, encompassing a wider range of licking and reward rates. We define a simple threshold for disengagement as licking bout rates below 1 bout/10 s, and reward rates below 1 reward/120 s. On average, mice are engaged 60.1% of the time. Figure 3B shows an illustrative example session in which the mouse transitions from task engagement to disengagement.

**Figure 3:**
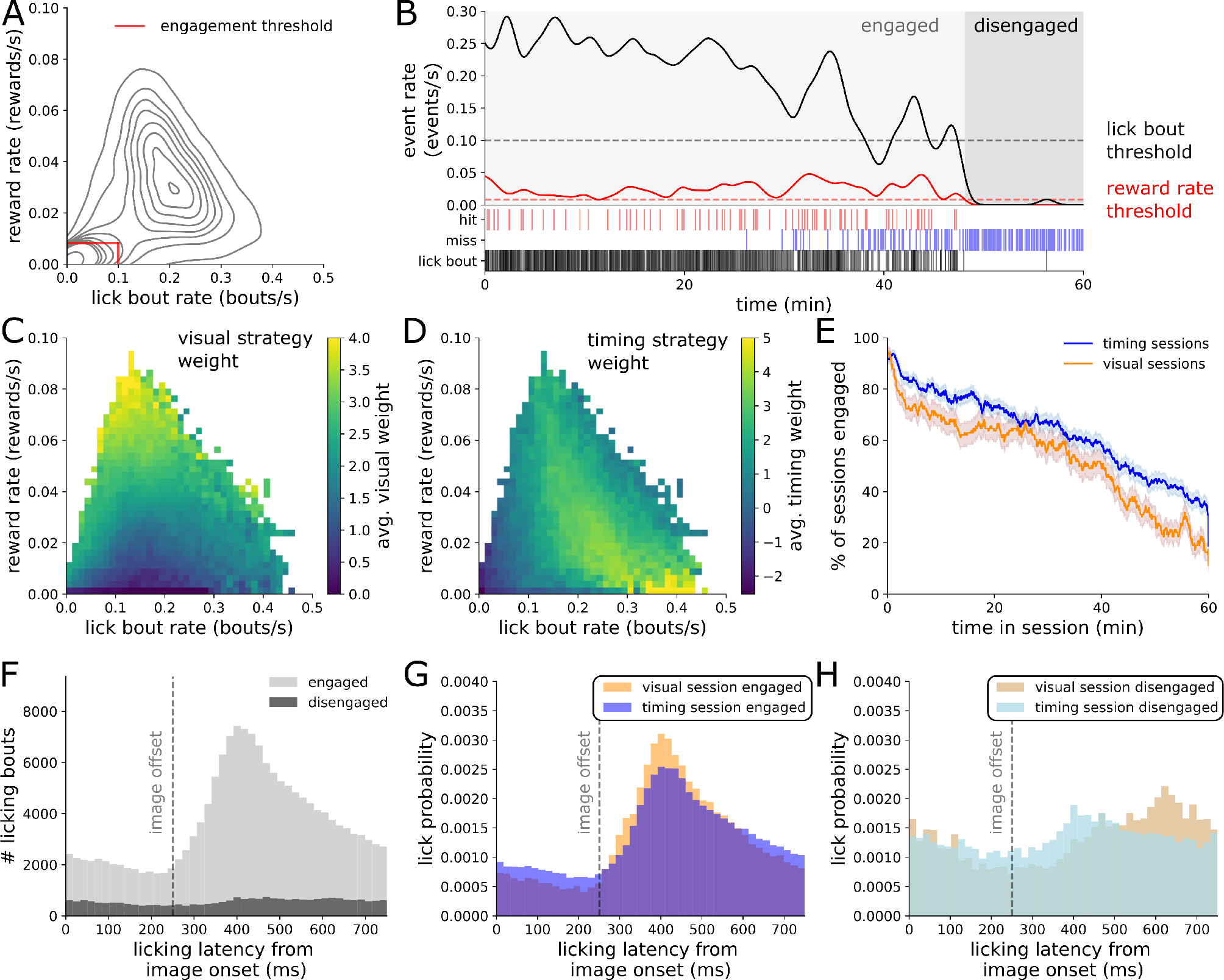
Strategy is distinct from engagement. (A) Contour plot of reward rate and lick bout rate from all imaging sessions (n=382 sessions, 1,804,462 image intervals). Red line marks our threshold for classifying engaged and disengaged behavior (1 reward/120s, 1 lick bout/10s). 60.1% of image intervals are classified as engaged. (B) Example session showing lick bout rate (solid black), licking engagement threshold (dashed black), reward rate (red), and reward rate threshold (dashed red). (C, D) Average value of the visual and timing strategy weights across a range of licking and reward rates. Both panels show data from all sessions (n=382 sessions, 1.8 million image intervals) across a range of licking and reward rates. (E) Percentage of sessions in an engaged state at each point in the hour long behavioral session, split by dominant strategy. (F) Response latency histogram split by engaged and disengaged epochs. Response latency is defined as the time from the start of each licking bout to the most recent image onset. (G) Response latency histogram for engaged periods, split by visual or timing strategy sessions. (H) Response latency histogram for disengaged periods, split by visual or timing strategy sessions.

In order to determine if task engagement was related to task strategy, we plotted the average strategy weights across the same landscape of licking bout and reward rates. Both the visual strategy weights (Fig. 3C) and timing strategy weights (Fig. 3D) are at their lowest values in the disengaged region. In the engaged region, the visual strategy is highest in the upper left when the ratio of rewards to licks is maximized. This can be understood as the visual strategy efficiently transforming licking bouts into rewards. The timing strategy is highest in the lower end of the engaged region, which can be understood as the timing strategy requiring more false alarm licks to generate rewards. From this analysis we conclude that task engagement is separate from each of the dominant strategies.

Next, we asked if the temporal pattern of engagement across each session was related to task strategy (Fig. 3E). For both strategies, engagement is highest at the start of the session and gradually decreases throughout the session, presumably as the mice become sated. A slightly higher percentage of timing strategy sessions are in the engaged state at each time point, presumably because the timing strategy requires more trial and error to earn water rewards.

Finally, we asked if the timing of licking was altered during disengaged periods by plotting histograms of when each licking bout started with respect to the latency since the last image presentation (Fig. 3F). By definition, the engaged periods have more licking bouts. However we see the disengaged licking bouts have altered timing relative to stimulus presentations. The engaged licking is time-locked to image onset, with the peak response time around 400ms. Disengaged licking lacks this clear time-locking to image onset. This pattern holds true for both behavioral strategies. Both visual and timing strategy sessions have clear time-locked responses in engaged epochs (Fig. 3G), and both lack clear time-locked responses in disengaged epochs (Fig. 3H). The fact that engaged licking in timing strategy sessions is time-locked to image presentations demonstrates that mice performing the timing strategy understand that rewards are tied to image presentations, and are synchronizing their timing-based guesses to image onsets rather than randomly licking without regard to the stimulus.

We conclude that mice performing both strategies go through periods of engaged and disengaged behavior, which is a separate dimension of behavior from their strategy preferences. Mice performing both strategies gradually disengaged over the course of the one hour behavioral session as they got sated by water rewards. Finally, mice performing both strategies have image-locked licking while engaged, and randomly timed licking when disengaged.

### Strategy is reflected in neural activity across the Vip-Sst microcircuit

We next wanted to assess whether the dominant behavioral strategy would be reflected in neural activity. To answer this question we turned to two-photon calcium imaging recordings during mouse behavior. We focused our analysis on two cortical visual areas, V1 and LM (Fig. 4A). Calcium imaging was performed in transgenic mice expressing the calcium indicator GCaMP6f in specific cell populations: excitatory neurons, Sst inhibitory neurons, and Vip inhibitory neurons. Recent surveys of neural cell types have proposed a taxonomy in which excitatory neurons and GABAergic neurons are classes, and Sst and Vip neurons are subclasses of GABAergic neurons (Tasic et al. 2018). In this paper, for simplicity we refer to excitatory, Sst, and Vip as cell classes. Each cell class was recorded in separate mouse populations. These three cell classes are thought to form a cortical microcircuit whereby Sst and Vip reciprocally inhibit each other (Fig. 4B). The dataset contains imaging while mice performed the task with both familiar and novel stimuli. Novel stimuli have dramatic effects on neural activity (Garrett et al. 2023), so we restricted our analysis to imaging during familiar image sessions. Our neural dataset contains 8,619 excitatory cells (21 imaging sessions, 9 mice), 470 Sst cells (15 imaging sessions, 6 mice), and 1,239 Vip cells (21 imaging sessions, 9 mice). We performed all of our analyses on discrete calcium events that were regressed from the raw fluorescence traces, thus removing the slow decay dynamics of the calcium indicator GCaMP6f (Fig. 4C). The extracted calcium events are of variable magnitude and correspond to a transient increase in internal calcium levels.

**Figure 4:**
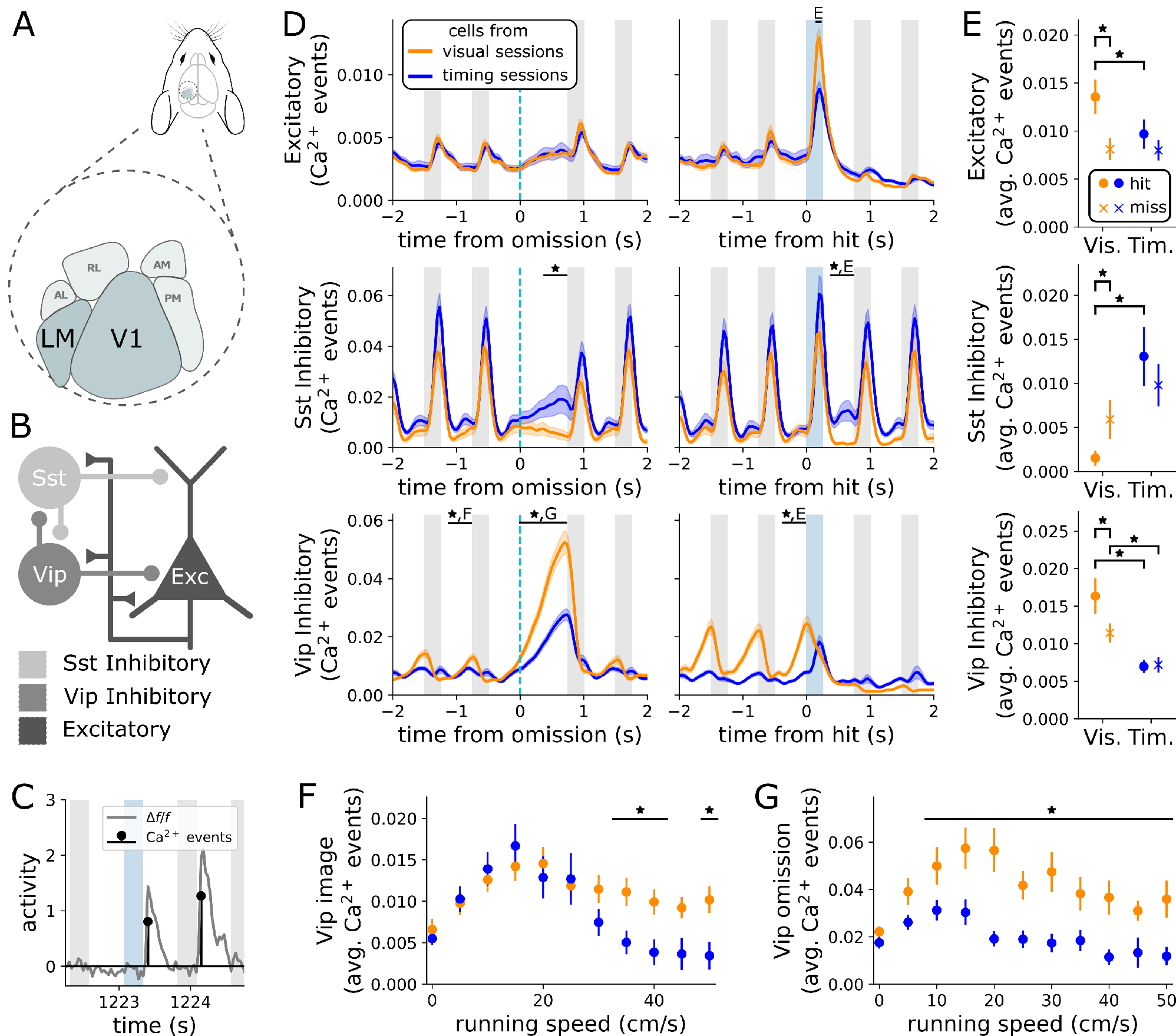
**Neural correlates of behavioral strategy across multiple cell populations**.(A) Two-photon calcium imaging was performed in visual areas V1 and LM. (B) Cartoon of VipSst microcircuit. Vip and Sst inhibitory neurons reciprocally inhibit each other. (C) Discrete calcium events were regressed from the fluorescence traces. (D) Average calcium event magnitude of each cell class aligned to image omissions (left column), and hits (right column), split by dominant behavioral strategy (* indicates p<0.05 from a hierarchical bootstrap over imaging planes and cells, corrected for multiple comparisons). (E) Average calcium event magnitude +/- hierarchically bootstrapped SEM in a interval around image changes split by strategy and whether the mouse responded. Excitatory and Sst cells show average events after image changes, (150, 250ms) and (375, 750ms) respectively. Vip cells show average events immediately before image changes (-375, 0 ms). * indicates p<0.05 from a hierarchical bootstrap over imaging planes and cells, corrected for multiple comparisons. (F) Average calcium event magnitude +/- hb. SEM in the 750ms interval after image presentations split by running speed and strategy (* indicate p<0.05 from a hierarchical bootstrap over imaging planes and cells, corrected for multiple comparisons). (G) Same as F after image omissions.

Our neural analysis examines how each cell class responds to three stimulus types: image repeats, distracting image omissions, and image changes (Fig. 4D). We grouped cells by the dominant strategy used by the mouse during the behavioral session in which they were recorded. We first asked how each cell class responds to image repeats, and if strategy affects responses to image repeats. Excitatory and Sst cells respond to each repeated image presentation with no difference in the population average between strategies. Excitatory cells are image selective, with each cell typically responding to only one of the 8 images presented during the session. In contrast Sst cells are broadly image tuned, thus the population average for excitatory cells is an order of magnitude smaller than Sst cells. Vip cells are suppressed by image presentations and ramp their activity between image presentations. Notably, Vip cells from visual strategy sessions showed significantly larger activity ramps between image presentations (Average calcium event magnitude +/- hierarchically bootstrapped SEM, visual 0.0096 +/- 0.00094, timing 0.0070 +/- 0.00092, p=0.025). We assessed significance and report the standard error of the mean (hb. SEM) by performing a hierarchical bootstrap (Saravanan et al. 2020). In summary, excitatory and Sst cells showed no strategy differences in response to image repeats, while Vip cells from visual strategy sessions showed increased activity between image repeats compared to cells from timing strategy sessions.

We next asked if strategy affects how each cell class responds to image omissions. In response to image omissions excitatory cells show an amplified response on the first image presentation after the omission, compared with the pre-omission image, with no difference in the population average between strategies. Sst cells showed significant strategy dependent changes in activity during the second half of the omission interval. Cells from timing strategy sessions show increasing activity ramping up over the omission interval, while cells from visual strategy sessions show decreasing activity during the omission interval (Average calcium event magnitude +/- hb. SEM, visual 0.0052 +/- 0.0021, timing 0.019 +/- 0.0054, p=0.0037). Sst cells from both strategies show decreased responses on the first image presentation after the omission compared to the pre-omission image. Vip cells show large ramps of activity during the omission interval with significant differences between strategies (Average calcium event magnitude +/- hb. SEM, visual 0.036 +/- 0.0038, timing 0.019 +/- 0.0022, p=0.00). In summary, Sst cells from visual strategy sessions show lower activity following omissions compared to cells from timing strategy sessions, while Vip cells from visual strategy sessions show increased ramping activity following omissions compared to timing strategy sessions.

We then asked if strategy affects how each cell class responds to image changes, including both hits (mouse licked) and misses (mouse did not lick). In response to image changes we see strategy dependent differences across all three cell classes. Figure 4E shows summary quantification of differences between hits and misses for each strategy.

We find excitatory cells from visual strategy sessions show greater activity in response to hits compared to cells from timing strategy sessions (Average calcium event magnitude +/- hb. SEM, visual hit 0.014 +/- 0.0017, timing hit 0.0097 +/- 0.0014, visual hit vs timing hit p=0.04). Further, we find a significant difference between hits and misses for cells from visual strategy sessions, but not for cells from timing strategy sessions (Average calcium event magnitude +/- hb. SEM, visual hit 0.014 +/- 0.0017, visual miss 0.0081 +/- 0.0010, timing hit 0.0097 +/- 0.0014, timing miss 0.0080 +/- 0.00095, visual hit vs visual miss p=0.0025). For Sst cells, cells from visual strategy sessions show lower activity during the interval after hits compared to cells from timing strategy sessions (Average calcium event magnitude +/- hb. SEM, visual hit 0.0015 +/- 0.00074, timing hit 0.013 +/- 0.0032, p=0.00). Further we find lower Sst activity following hits compared to misses for Sst cells from visual but not from timing strategy sessions (Average calcium event magnitude +/- hb. SEM, visual hit 0.0015 +/- 0.00074, visual miss 0.0059 +/- 0.0021, timing hit 0.013 +/- 0.0032, timing miss 0.0098 +/- 0.0022, visual hit vs visual miss p=0.010). For Vip cells, in the interval before the image change we find significant differences in activation between the two strategies as well as between hits and misses for cells from visual but not from timing strategy sessions (Average calcium event magnitude +/- hb. SEM, visual hit 0.016 +/- 0.0022, visual miss 0.011 +/- 0.0011, timing hit 0.0070 +/- 0.00076, timing miss 0.0072 +/- 0.00087, visual hit vs visual miss p=0.037, visual hit vs timing hit p=0.00, visual miss vs timing miss p=0.0003). In summary, cells from visual strategy sessions show differential activity in all three cell classes between hits and misses, while cells from the timing strategy shows no modulation between hits and misses. Further, we see strategy dependent differences in neural activity in responses to hits across all three cell types: excitatory cells from visual strategy sessions show greater activity after hits compared to cells from timing strategy sessions, Sst cells from visual strategy sessions show lower activity following hits compared to cells from timing strategy sessions, and Vip cells from visual strategy sessions show greater activity before hits and misses compared to cells from timing strategy sessions.

Vip cells are known to be modulated by locomotion (Fu et al. 2014). This effect is seen in our data most visibly in figure 4D when the mice stop running after image changes to consume their water reward. With this in mind, we wanted to know whether the strategy differences we observe could be due to differences in running speed or patterns. To answer this we looked at the Vip response amplitude to images and omissions as a function of running speed (Fig. 4F,G). Broadly, we observe Vip activity in cells from visual strategy sessions is equal to or greater than cells from timing strategy sessions across all running speeds. From this we conclude that strategy differences in Vip cell activity cannot be a result of different running speeds or patterns.

### Microcircuit disinhibition dynamics are amplified in the visual strategy

In order to make sense of the diverse strategy differences we observe in figure 4 we now turn to the Vip-Sst microcircuit as a unifying model (Fig. 5A). We first asked how the microcircuit components respond as a system to each stimulus type. We thus examined the neural population averages grouped by strategy allowing a direct comparison across cell classes (Fig. 5B). We emphasize here each cell class was recorded in a separate population of mice. Viewing the neural population averages in this manner highlights that excitatory and Sst cells are image responsive, while Vip cells are inhibited by image presentations. Notably Vip cells show ramping between image presentations. We can further condense this information by showing the three cell classes in a 3D state space plot (Fig. 5C), or for clarity in a 2D state space between excitatory and Vip cells. These state space plots reveal the dynamics of the three cell classes as a periodic cycle corresponding to the rhythmic stimulus presentations where the two strategies differ primarily in the Vip activation between image presentations. Importantly, in this periodic cycle excitatory and Sst cells are tightly correlated, each responding to image presentations, while Vip cells are suppressed by image presentations. Supplemental Note 9 shows 3D state space plots and all 2D state space combinations of cell classes. We will next consider image omissions and image changes as perturbations to this underlying microcircuit cycle.

**Figure 5:**
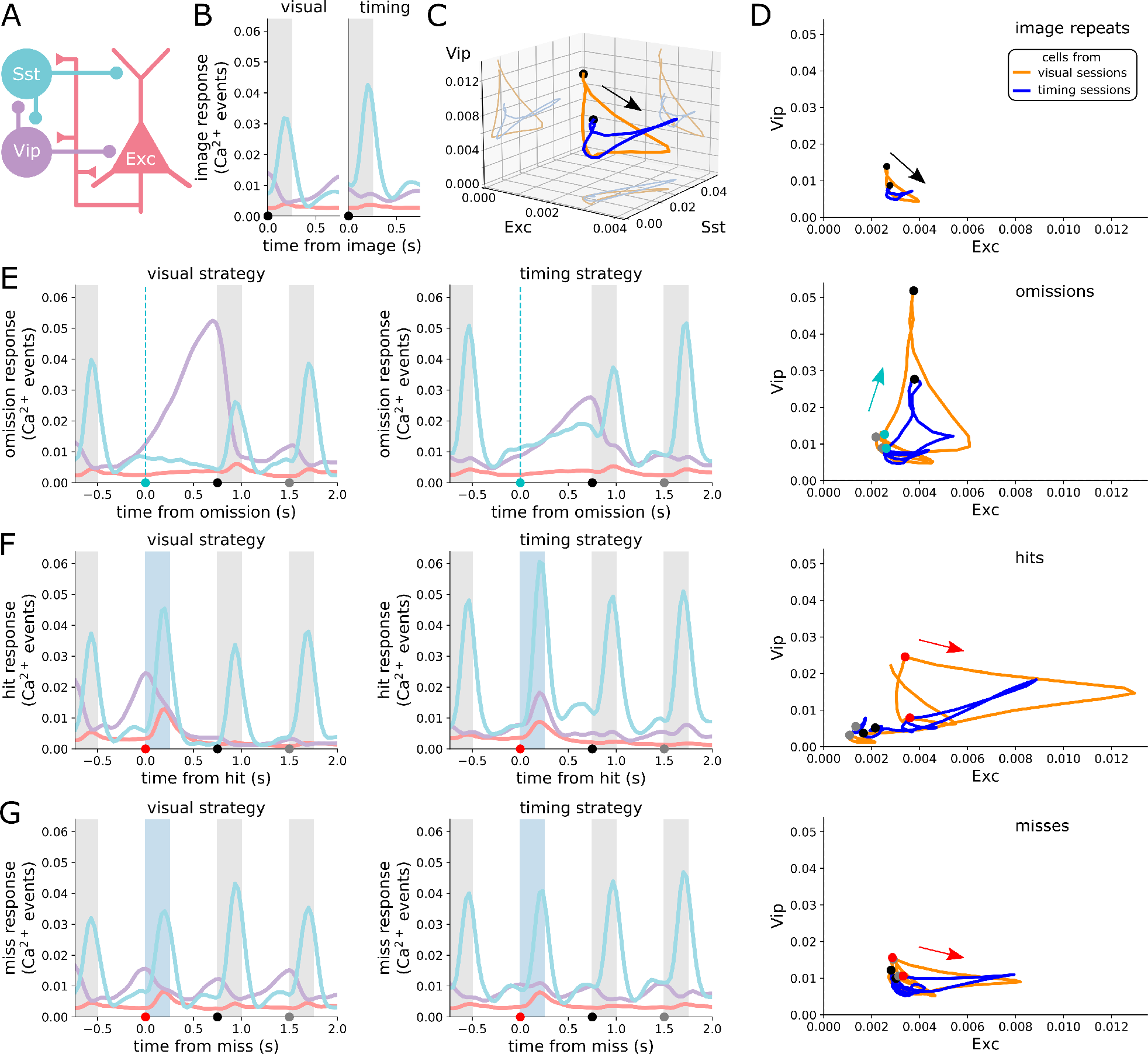
Microcircuit disinhibition dynamics are amplified in the visual strategy (A) Cartoon schematic of microcircuit. (B) Population average response to image repeats, grouped by strategy. (C) Population average response to image repeats plotted in 3D space. (D) Population average response to image repeats for excitatory cells against Vip cells. (C,D) Arrow marks forward progression of time. Black circle marks image onset in B. (E) Same as B,D for image omissions. (F) Same as B,D for hits. (G) Same as B,D for misses.

Previous work (Millman et al. 2020) found that Vip-Sst antagonism regulates the gain of cortical circuits to broaden the range of stimulus contrast levels excitatory cells can encode. Comparing the dynamics of Vip and Sst cells in response to image omissions we can see this antagonism at work (Fig. 5E). When an image is omitted, the Vip cells continue their between-image ramping until they are finally suppressed by the post-omission image. Sst cells have smaller responses to the post-omission image compared to the pre-omission image and excitatory cells have larger responses to the post-omission image compared to the pre-omission image, presumably a result of the increased Vip activation. We can interpret this ramping as Vip cells responding to the lack of visual stimulus by amplifying the gain of the cortical circuit via inhibition of Sst cells and disinhibition of excitatory cells. In the state space view of microcircuit dynamics, the image omission is a perturbation to the periodic repeating image cycle. In response to this perturbation, Vip cells activate, releasing excitatory cells from inhibition thereby amplifying the response to the post-omission image before returning to the periodic image cycle on subsequent images. Notably, these gain dynamics are amplified in cells from visual strategy sessions compared to cells from timing strategy sessions (Fig. 5E).

Image changes can also be understood as perturbations to the periodic image cycle. In response to both hits and misses excitatory cells show increased responses. For cells from visual strategy sessions we see notable differences between hits and misses in Vip activity before the image change (Fig. 5F,G). We do not observe these differences for cells from timing strategy sessions (Fig. 4E). One possible interpretation of these data is that mice performing the visual strategy are more likely to make the choice to respond or not based on visual cortex activity. On a trial by trial basis, a more active Vip population may prime excitatory cells for more robust change responses. Thus leading to increased Vip activity before hits compared to misses, increased excitatory activity on hits compared to misses, and decreased Sst activity after hits compared to misses. We emphasize here that the increased excitatory response on hits happens, on average, 200ms before the mouse responds and therefore cannot be interpreted as a reward response. In this interpretation, mice performing the timing strategy are less reliant on visual cortex activity to trigger responses, instead using some internal timing mechanism elsewhere in the brain. Thus, we see no differences in average Vip, Sst, or excitatory activity between hits and misses for timing mice. Increased Vip activity priming visual, but not timing, strategy mice to respond may also explain why visual strategy mice are more likely to respond to the post-omission image (Fig. 2E). In the state space view of microcircuit dynamics, the image change perturbs the ongoing periodic image cycle by elevating excitatory responses. For mice performing the visual strategy, but not the timing strategy, increased Vip activity can amplify excitatory responses and preferentially lead to behavioral responses.

In summary, when comparing between visual and timing strategy circuits, we can succinctly describe the differences in population response to image repeats, omissions, and changes as amplified Vip disinhibition dynamics. Excitatory and Sst cells respond in unison to repeating image presentations, while Vip cells are suppressed by each image presentation, instead ramping their activity between images. This between image ramping is elevated in cells from visual strategy sessions. When images are omitted, Vip cells continue to ramp their activity which in turn suppresses Sst and disinhibits excitatory cells during the post-omission image. This functionally allows Vip cells to calibrate the subsequent gain of excitatory cells, and this mechanism is heightened in cells from visual strategy sessions. Finally, for the visual strategy elevated Vip activity before image changes facilitates increased excitatory responses which in turn potentially leads to mouse licking responses.

### Trial by trial neural activity is more correlated with behavioral choices for visual strategy mice

Following the interpretation suggested above that visual strategy mice are more reliant on visual cortical activity to drive choices, we next wanted to determine if the differences in neural activity between strategies were behaviorally relevant on an trial by trial basis. To answer this we used a random forest classifier trained on neural activity to predict either image changes versus repeats (change decoder), or hits versus misses (hit decoder). Decoding was performed on neural activity in the first 400ms after each stimulus presentation. This time window is before the average licking response, and thus largely avoids reward signals.

Across all three cell types, we found that image changes versus repeats could be decoded equally well from cells from visual strategy sessions and timing strategy sessions (Fig. 6A). We then asked to what degree the image change signals were correlated with animal behavior. Since we observe differential activity on hits versus misses for cells from visual strategy sessions, but not cells from timing strategy sessions, we reasoned that change signals in visual cortex should be more correlated with behavior when mice are performing the visual strategy. We measured the correlation between the decoder’s predictions on image changes (change vs repeat) and the animal’s choice (hit vs miss). For excitatory, but not Vip or Sst cells, we find a stronger correlation for cells taken from visual strategy sessions compared to timing strategy sessions (Fig. 6B). For excitatory cells, the correlation between behavior and decoder predictions is more than twice as strong for visual sessions compared to timing sessions. This demonstrates that while image change information is equally present in neural activity from mice performing both strategies, it is more correlated with animal choices in visual strategy sessions. This finding is consistent with the interpretation that mice performing the visual strategy are more dependent on activity in visual cortex to drive behavioral responses. Finally, we asked how well we could decode hits versus misses. For excitatory and Vip cells, but not Sst cells, decoder performance was higher for visual strategy sessions compared with timing strategy sessions.

**Figure 6:**
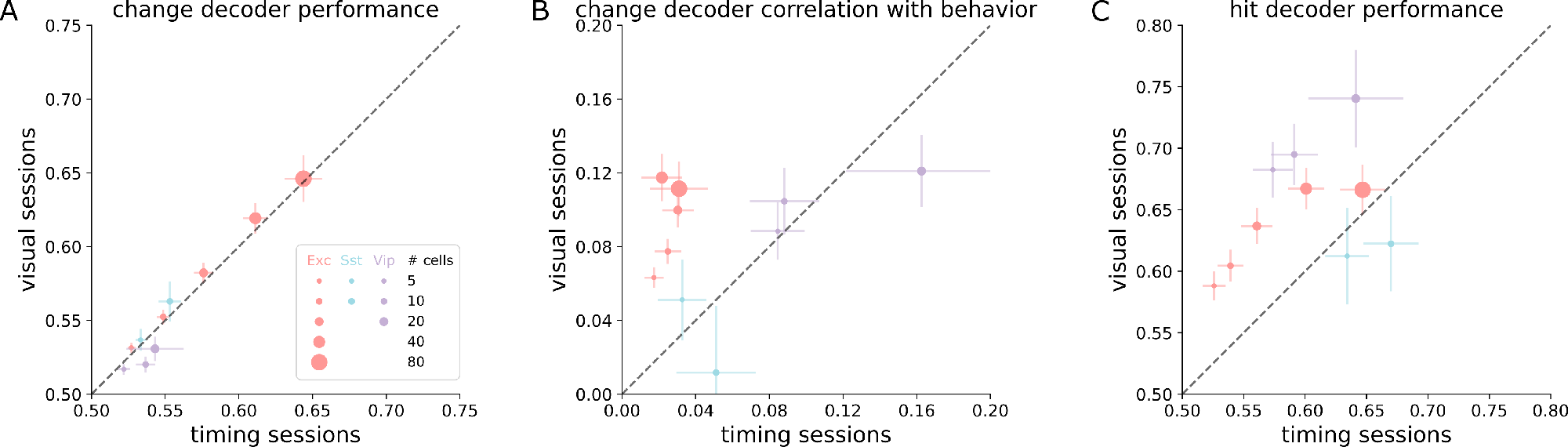
Stronger behavioral choice signals in cells from visual strategy sessions. (A-C). Decoding was performed on neural activity in the first 400ms after image presentation. Error bars are SEM over imaging planes. Each cell type is plotted as a separate color, with marker size indicating the number of cells used for decoding from each imaging plane. (A) Cross validated random forest classifier performance at decoding image changes and repeats (% correct). (B) Correlation between decoder prediction on image changes (change vs repeat) and animal behavior (hit vs miss). (C) Cross validated random forest classifier performance at decoding hits and misses (% correct).

### Engagement state has limited modulation of neural activity

We next asked if we could observe neural correlates of task engagement. Similar to figure 4D, we plotted the average neural activity aligned to either image omissions or image changes but now additionally split by engaged and disengaged states (Fig. 7). The disengaged state does not contain hits, so we restricted our comparison to misses. Excitatory cells from visual strategy sessions, but not timing strategy sessions, show elevated responses to misses when engaged compared to disengaged (Average calcium event magnitude +/- hb. SEM, visual engaged 0.010 +/- 0.0016, visual disengaged 0.0069 +/- 0.00088, p = 0.032). We do not observe any other significant differences between average neural activity in engaged and disengaged states for either dominant strategy across any cell type. Further, we do not see major differences in Vip activity even when controlling for running speed differences during engagement periods (Supplemental Note 10). Thus, we conclude that the effects of strategy are separate from task engagement.

**Figure 7:**
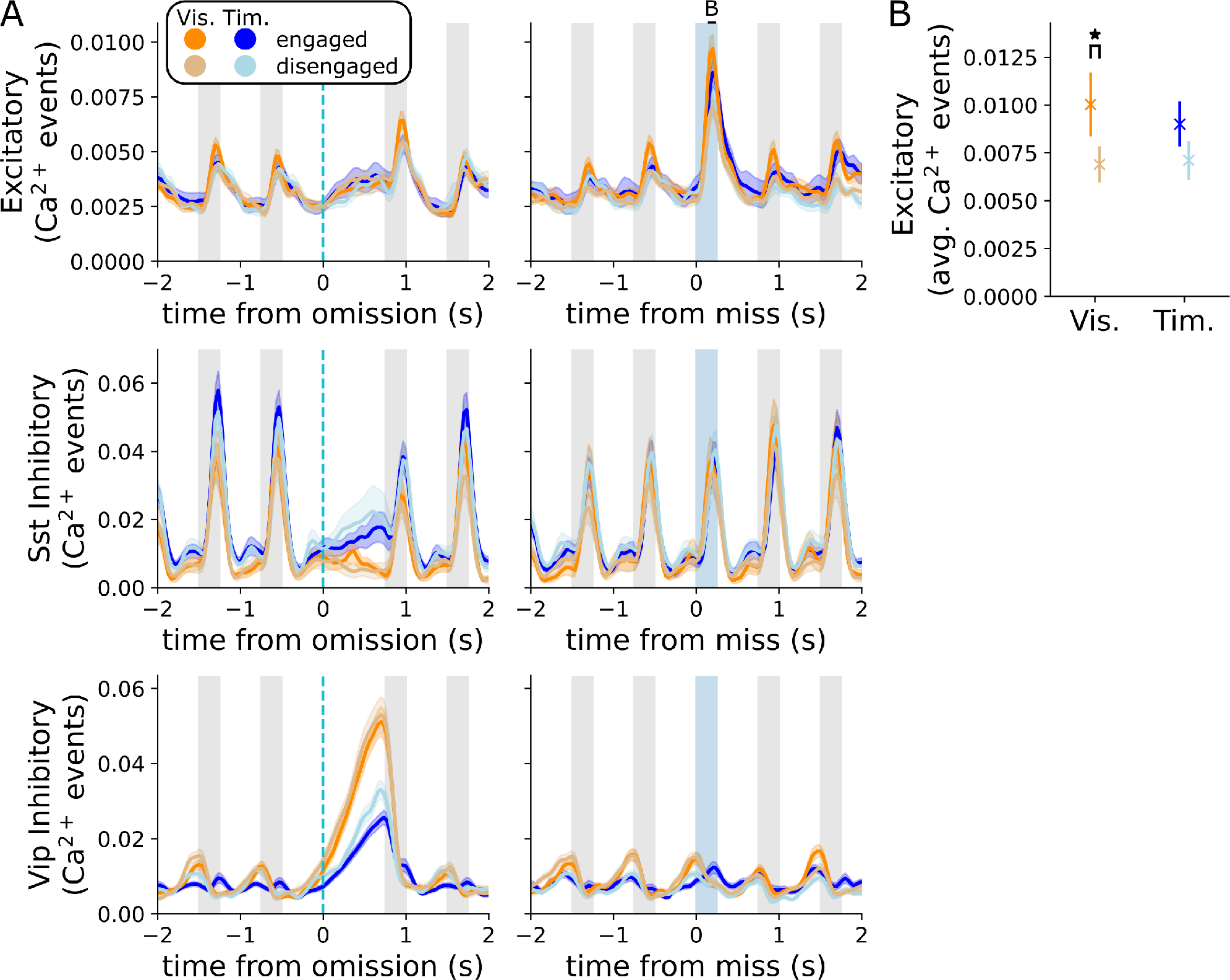
**Task engagement has minor effects on neural population activity**. (A) The average activity of cells from each dominant task strategy, split by epochs of task engagement and disengagement, aligned to either image omissions (left), or image change misses (right). Error bars are +/- SEM. (B) Average calcium event magnitude +/- hierarchically bootstrapped SEM in a interval (150ms, 250ms) around image changes split by strategy and whether the mouse responded. * indicates p<0.05 from a hierarchical bootstrap over imaging planes and cells, corrected for multiple comparisons.

### Both dominant strategies show robust effects of image novelty

Our analysis to this point has examined neural activity when mice are shown familiar stimuli they have seen many times. The Allen Institute Visual Behavior - 2p calcium imaging dataset also contains neural activity in response to novel stimuli. Garrett et al. 2023 examined this data and found exposure to novel stimuli dramatically altered neural activity, finding striking effects across all three recorded cell classes. We sought to extend the results reported in Garrett et al. 2023 by asking two questions. First, behaviorally, does strategy change with novel stimuli? Second, how do cells from each dominant strategy respond to novel stimuli? Supplemental Note 11 answers both questions. Analyzing behavior, we see a small but significant shift in strategy preference towards the visual strategy on the novel image session. This shift manifests as most mice slightly increasing their strategy index, rather than individual mice making dramatic changes in strategy. Analyzing neural activity we see cells from both strategies show the effects of novelty documented in Garrett et al. 2023. Further, on the novel session the primary effects reported in figure 4 are still present. Namely, Vip cells from visual strategy sessions had increased activity on image omissions, and before hits compared to cells from timing strategy sessions. As well as increased excitatory activity for hits compared to misses for cells from visual strategy sessions but not from timing strategy sessions. We conclude the effects of stimulus novelty are largely separate from strategy preference.

## Discussion

The Allen Institute Visual Behavior - 2p calcium imaging dataset (brain-map.org) is a large scale survey of neural activity in three neural cell classes in the context of a change detection task. We sought to identify behavioral strategies used by mice in this dataset. Using a dynamic logistic regression model we found that mice have unique mixtures of two strategies, a visual comparison strategy where mice respond to image changes, and a statistical timing strategy where mice respond at the expected duration between image changes. Individual mouse strategy preferences were relatively stable over multiple behavioral sessions and emerged gradually over training. Separately from strategy preference, we found mice have periods of task engagement and disengagement. Mice performing either strategy gradually disengaged from the task throughout each behavioral session. For both dominant strategies, engaged licking was time-locked to stimulus presentations, while disengaged licking was not time-locked to stimulus presentations.

We then analyzed neural activity based on the dominant strategy preference. We found a diversity of neural correlates of strategy across excitatory, Sst inhibitory, and Vip inhibitory cell classes. We found that this diversity of responses can be understood through the lens of the Vip-Sst disinhibitory circuit. In response to image omissions, Vip cells increase their activity to change the gain on excitatory cells. In visual strategy sessions, this Vip gain dynamic is amplified. Further, in visual strategy sessions, but not timing strategy sessions, Vip activity is elevated before hits compared to misses, suggesting Vip cells potentiate excitatory cells and bias the decision to respond to each image presentation in mice performing the visual strategy. Supporting the view that visual strategy mice are more reliant on visual cortical activity, we found stronger hit decoding from visual strategy sessions, and the performance of an image change decoder is more correlated with animal behavior in visual strategy sessions. Additionally, the visual strategy mice show increased post-omission licking compared to timing strategy mice, suggesting Vip omission activity potentiates excitatory responses on the post-omission image.

Despite clear effects of task strategy, we found only limited neural correlates of task engagement. Excitatory cells from visual strategy sessions show elevated responses to misses when engaged. This suggests the effects of strategy influence neural circuits on longer timescales. Additionally, we found the effects of strategy preference persist when the mice perform the task with novel stimuli, despite robust changes to population activity. Our findings demonstrate that behavioral strategy alters neural activity within visual cortex, and is mediated by specific cell classes.

### Behavioral diversity

We note that it could have been possible to alter our training pipeline to push mice away from the timing strategy, skip over mice performing the timing strategy for neural imaging, or posthoc exclude their data from neural analysis. Indeed, such practices are common in neuroscience laboratories where strict experimental control is demanded by the practicalities of limited experimental resources and a desire to clearly isolate single behaviors or computations. Recently there has been considerable discussion over the advantages of naturalistic behavior and tightly controlled laboratory tasks (Juavinett et al. 2018; Musall et al. 2019). Naturalistic behavior offers ethological relevance and behavioral richness, while laboratory tasks can isolate behaviors and yield reproducibility. Our findings demonstrate a middle ground, through the use of large scale brain observatories with a behavioral task that subjects can solve in multiple ways. By recording from many mice we find considerable behavioral richness across our population, but still harness the advantages of well defined stimuli and task structure. This approach compliments the existing range of behavioral paradigms in the field.

### How do behavioral strategies arise?

We found that strategy preferences emerge gradually over many days of training. How do mice learn their strategy preferences? One possibility is mice could adopt one strategy or another based on a myriad subtle biases such as visual ability, cognitive ability, or difficulty licking the reward spout. Alternatively, mice could have been biased to one strategy based on variability in early behavioral exploration. Given that the visual strategy can earn a greater number of rewards per session we can consider the timing strategy as a local maximum in behavioral performance that some mice might settle into early in training. Finally, it is possible that prior to any task training, some mice could already have amplified Vip-Sst dynamics and thus be predispositioned to adopting the visual strategy. This predisposition could be the result of genetics, or life experience. The learning processes that lead mice to each strategy may involve unique behavioral states and neural mechanisms (Rosenberg et al. 2021; Meister 2022).

### How does strategy preference amplify Vip-Sst circuitry?

We found that the visual strategy resulted in amplified Vip-Sst disinhibition dynamics. If strategy preferences arise from outside visual cortex, then what mechanisms facilitate the diversity of neural correlates we observe? Vip cells are known to be preferential targets of top-down feedback (Kamigaki 2019; Ma et al. 2021). One possibility is that external feedback results in higher tonic activation of Vip cells during visual strategy sessions. Alternatively, neuromodulatory input could alter intrinsic Vip firing patterns (Férézou et al. 2002; Fu et al. 2014; Prönneke et al. 2020).

Strategy preference could also recruit different visual pathways and brain structures (Bolkan et al. 2022). We found cells from visual strategy sessions, but not timing strategy sessions, show differential activity between hits and misses. Further, we found a decoder trained to predict image changes made predictions that were more correlated with the animal’s choices for visual sessions compared to timing sessions. For excitatory cells, the correlation was more than twice as strong. These results suggest that the timing mice may execute this task primarily through other brain structures. We re-iterate here that licking bouts from timing mice are time-locked to image presentations. This demonstrates timing strategy mice still use visual input, but perform less processing of the visual stimulus. Retinal ganglion cells project to many sub-cortical structures (Martersteck et al. 2017), which could facilitate the simpler visual processing required for the timing strategy.

### Engagement and novelty

Mice performing both dominant strategies displayed periods of task engagement and disengagement (Ashwood et al. 2022). Across both dominant strategies we observed limited modulation of population activity with task engagement. This may be a puzzling finding, especially given some similarities between task engagement and visual attention. However, we caution that task engagement is a separate phenomena from visual attention and care should be taken before taking inspiration from attentional mechanisms. With this reservation in mind, we do note that Myers-Joseph et al. 2023 found that attention modulation operates distinctly from Vip disinhibition. Consistent with our findings, Pho et al. 2018 found little modulation of neural activity by engagement in V1 but significant modulation in posterior parietal cortex.

One possibility for a lack of engagement correlates is the nature of our stimuli. Our stimuli are large full visual field images with relatively high contrast they are not near perceptual thresholds. Perhaps our stimuli are salient enough to evoke visual responses regardless of task engagement. If our task operated at a perceptual threshold we might observe stronger modulation of neural activity by task engagement.

The Allen Institute Visual Behavior - 2p calcium imaging dataset contains multiple behavioral dimensions, including novel stimuli. We did not focus on novel stimuli, but Garrett et al. 2023 found striking changes in neural activity in all three recorded cell classes. We found strategy alters neural activity in distinct ways from novel stimuli. Since strategy differences are present during both task engaged and disengaged states, it is likely the strategy effects we observe are the result of long-term learning and firmly established in cortical circuits. This is further supported by the persistence of strategy differences during the novel stimulus presentation. The neural correlates of strategy on this task are therefore not a fleeting activity pattern, or a flexible state to be switched on and off, rather it appears to be a deeply ingrained change in the cortical circuit that the mouse develops to solve the task consistently well.

### Future directions

A naive view of sensory circuits might expect veridical encoding of the sensory world. However our findings highlight that task strategy is mediated through changes in neural activity in sensory cortex. This result raises general questions about how, when, and why cognitive states alter sensory processing. Future studies should seek mechanistic understanding of how cognitive states influence local circuit processing in visual cortex, how local circuit processing changes with learning, and how cognitive states influence the propagation of sensory information up the visual hierarchy into deeper brain structures.

## Methods

### Data selection

The collection and processing of all data in the study was previously described in Garrett et al. 2023, and is available at https://portal.brain-map.org/explore/circuits/visual-beh avior-2p. For our behavioral analysis we used all active behavioral sessions from mice in the V1 and LM datasets across all image set experience levels (familiar images, novel images, and repeated exposure to novel images). For all analyses we combined mice trained on image sets A and B. For neural analysis we used neurons recorded during familiar image set presentations on the multi-plane imaging rig. Except where noted we combined across cells from V1 and LM and across cortical depths.

### Behavioral data processing

We performed all of our behavioral analysis after assigning behavioral events to each image presentation interval. By image presentation interval we refer to the 750ms interval beginning with each image presentation. For image omissions we used the 750ms following the time of the omission, when the image should have been presented. Licks were segmented into licking bouts using an inter-lick interval of 700ms. This threshold was determined by visual inspection of the histogram of inter-lick intervals. The start and end of each licking bout was then assigned to an image presentation interval.

### Strategy model

The strategy model predicts whether a licking bout started on each image presentation interval. We thus excluded from our model fits any image presentation intervals where a licking bout was on-going at the start of the interval. The licking bias strategy vector was defined as a 1 on every interval. The visual, omission, and post-omissions strategy vectors were defined as 1 on intervals with the respective stimulus and 0 otherwise. The timing strategy vector was a number between 0 and 1 based on a sigmoidal function of how many image intervals since the end of the last licking bout. See supplemental note 3 for details on the timing strategy regressor. Except for the licking bias, all strategy vectors where then mean-centered.

### Fitting the strategy model

We fit the strategy model using the PsyTrack package (Roy et al. 2018; Roy et al. 2021, https://github.com/nicholas-roy/psytrack). The PsyTrack package fits the model through an empirical Bayes procedure. The hyper-parameters are first selected by maximizing the model evidence. Then the strategy weights were determined by the MAP estimate. The model hyperparameters and strategy weights were fit separately for every behavioral session.

Model performance was determined using the cross-validated model predictions as a classifier to determine whether a mouse initiated a licking bout on each image interval. The receiver operator curve determines the rate of true positives against false positives as a function of different classifier thresholds. The area under the curve provides a summary statistic to compare models. Behavioral sessions were classified as visually dominant or timing dominant by determining which strategy lead to the greater decrease in model evidence when that strategy was removed.

### Task Engagement

Task engagement periods were determined by applying a threshold to the reward rate and lick bout rate. Task engagement was determined for each image presentation interval. Both rates were calculated by annotating which image presentations intervals had rewards and lick bout initiation. We then smoothed across image presentations with a filter. We used a Gaussian filter with standard deviation of 60 images. We then converted both rates into units of events per second. We set the thresholds of 1 reward per 120 seconds and 1 lick bout per 10 seconds through visual inspection of the behavioral landscape in figure 3A. If either rate was above its threshold, then the interval was labeled engaged.

### Neural data

For all analysis of neural data we used the detected calcium events as described in Garrett et al. 2023 and https://portal.brain-map.org/explore/circuits/visual-behavior-2p. This process produces, for each cell, a set of calcium events each with a time and magnitude. Except where noted we used familiar image set sessions collected on the multiplane calcium imaging rig. To generate the population averages traces we compute the behavioral event triggered response for each cell to each behavioral event and then average across all cells in each population. We compute the behavioral event triggered response by isolating the calcium events around the triggering behavioral event, then linearly interpolating onto a consistent set of 30hz timestamps relative to the triggering behavioral event. This produces a vector of calcium event magnitudes for each cell relative to each behavioral event (omission, image change, or repeat).

To generate the average calcium response across a time interval, we first compute the event triggered response for each cell as described above. We then average across all time points in the same image interval, producing a single scalar for each cell on each image interval. Average calcium responses were computed for either the entire image interval (50, 800ms), the first half of each image interval (50, 425ms), the second half of each image interval (425, 800ms), or a more narrow stimulus locked window for excitatory cells (150, 250ms). We used a 50ms delay for the image interval, (50, 800ms) rather than (0, 750ms), to account for signal propagation to visual cortex.

### Hierarchical bootstrap analysis

To determine significance for average calcium response metrics we applied a hierarchical bootstrap method (Saravanan et al. 2020). On each bootstrap iteration we sampled with replacement from first imaging planes, and then for each imaging plane we sampled with replacement from cells from that plane. Averaging across all of these cells produces one bootstrap sample. For all of our analyses, we used this procedure to generate 10,000 samples. The standard deviation of this set of samples produces an estimate of the standard error of the mean. To performance hypothesis testing we assigned samples from each condition into random pairs and performed pairwise comparisons to determine what fraction of samples from each condition was greater or less than the other condition. In this context an imaging plane is a specific cortical area and depth from one behavioral session, so by sampling imaging planes we are effectively sampling over sessions and mice. We corrected for multiple comparisons through the Benjamini-Hochberg procedure (Benjamini et al. 2001).

### Running speed

Running speed traces were processed in the same manner as calcium event traces. For each behavioral session, we computed the event triggered running trace by isolating running timepoints around the triggering behavioral event then linearly interpolating onto a common 30hz timeseries. We then averaged across all points in the relevant time window.

### Decoding analysis

To decode task signals on an image by image basis we used a random forest classifier to predict either image changes versus repeats (change decoder), or hits versus misses (hit decoder). We iterated over the number, n, of simultaneously recorded neurons used in the decoding analysis. For each imaging plane we sampled neurons and performed decoding until there was a 99% probability all neurons had been used in decoding. For each sample we took n neurons without replacement and concatenated their neural activity on each image presentation to make a k *×* nt matrix X. Here k is the number of images, and t is the number of timesteps from each neuron on each image. The change decoder used each image change and the image repeat immediately before the image change. The hit decoder used all image changes. We then performed 5-fold cross validated decoding using the RandomForestClassifier package from *sklearn*. We evaluated decoder performance as the percentage of test-set images correctly classified. We averaged the performance of all samples from the same imaging plane and report summary statistics as the mean +/- SEM over imaging planes. We computed the correlation between the change decoder’s predictions and the animal’s choices using the phi coefficient.

## Acknowledgements

We thank the Allen Institute founder, Paul G. Allen, for his vision, encouragement, and support. We thank the members of Allen Institute for a fruitful scientific community and helpful discussions. AP thanks Tyler Boyd-Meredith for comments on the manuscript, and Nick Roy for developing the dynamic logistic regression model, the associated code package PsyTrack, and helpful discussions.

## Author contributions

AP conceived of the project, performed behavioral and neural analysis, and wrote the paper. NP and DO performed behavioral analysis. DO, MG, PG, SO, and CK oversaw task development and data collection. AA oversaw the project.

## Data availability

All data used in this study is publicly available at https://portal.brain-map.org/explore/circuits/visual-behavior-2p

## Code availability

The behavioral analysis code is available at https://github.com/alexpiet/licking behav ior/. The strategy model fits are available at https://figshare.com/projects/Allen Ins titute Visual Behavior Strategy Paper/160972. The neural analysis code is available at https://github.com/AllenInstitute/visual behavior glm.

## Supplemental Materials

The supplementary materials contains extended figures and control analyses for several aspects of the study:

1. Mouse behavior
2. Segmenting licks into licking bouts
3. Constructing the timing strategy
4. Model validation
5. Strategy characterization
6. PCA on model strategies
7. Strategy over training
8. Strategy behavior over time
9. Microcircuit dynamics
10. Running speed and task engagement
11. Transition to the novel image set

## Supplemental Note 1 - Mouse behavior

**Figure 8:**
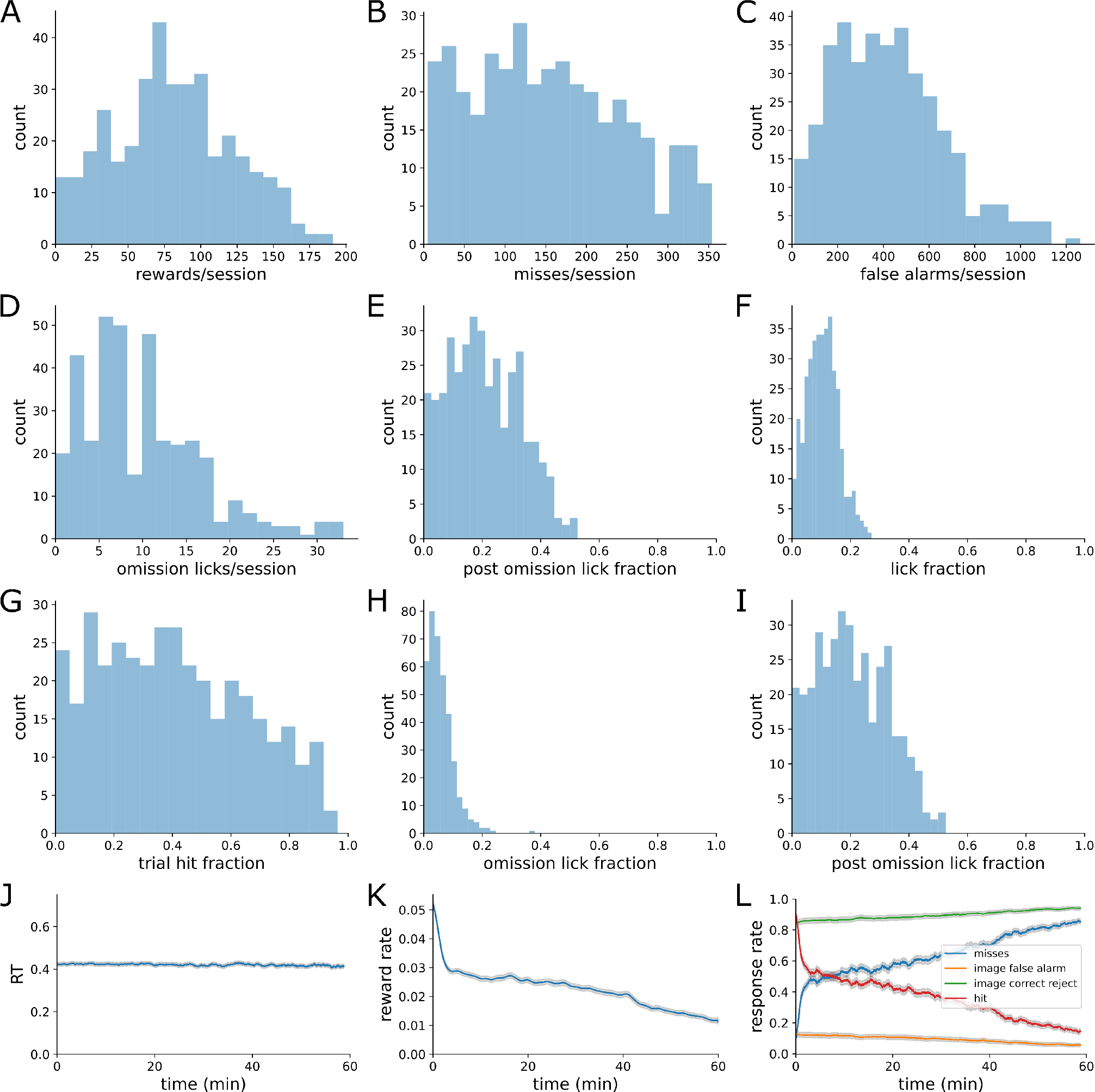
Quantification of mouse behavior. (A) Histogram of rewards/session. (B) Histogram of misses/session. (C) Histogram of false alarms / session. (D) Histogram of omissions with licks/session. (E) Histogram of post-omission-licks/session (F) Histogram of average lick fraction per session. (G) Histogram of fraction of image changes with licks per session. (H) Histogram of fraction of omissions with licks per session. (I) Histogram of fraction of post-omission images with licks per session. (J) Average Response latency over time. (K) Average reward rate over time. (L) Hit, Miss, False alarm, and correct reject rates over time.

## Supplemental Note 2 - Segmenting licks into licking bouts

**Figure 9:**
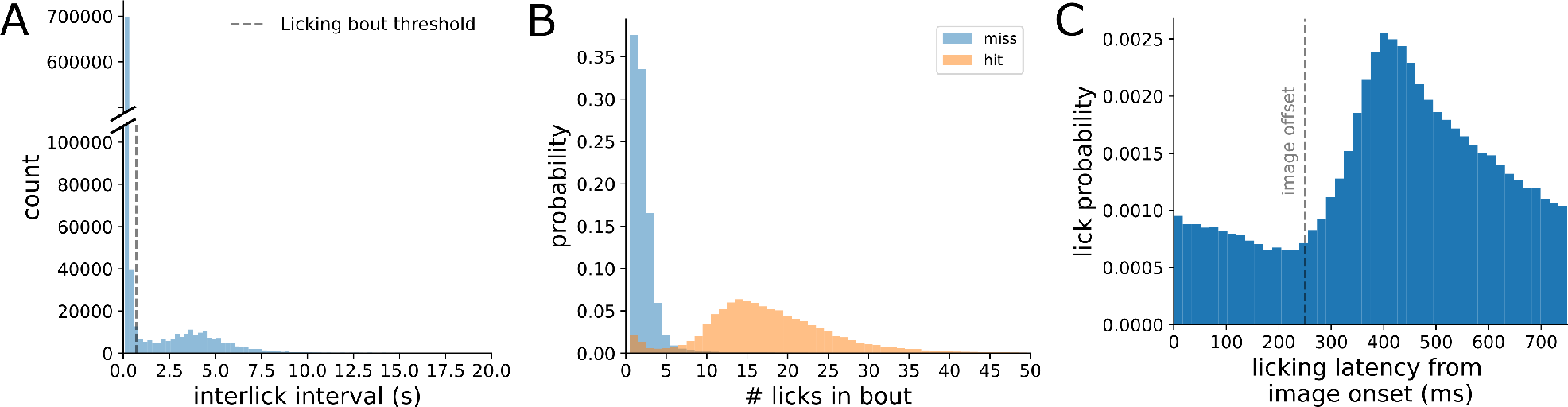
Licks were segmented into licking bouts and aligned to image onset. (A) Histogram of interval between successive licks (n = 936,136 licks from 382 imaging sessions). Dashed line indicates 700ms threshold used to separate licks within the same licking bout (< 700ms) and licks in separate licking bout (> 700ms). (B) Histogram of the number of licks in each licking bout separated by whether the licking bout earned a reward (hit) or did not (miss) (n = 190,410 licking bouts from 382 imaging sessions). (C) Histogram of the response latency for the start of each licking bout with respect to the most recent image onset (n = 190,410 licking bouts from 382 imaging sessions).

## Supplemental Note 3 - Constructing the timing strategy

A subset of 45 sessions were used to construct the timing regressor. The strategy model was fit with 10 timing regressors, each using 1-hot encoding for different length delays since the image with the end of the last licking bout. Then a four parameter sigmoid was fit to the average weights of each of these 1-hot timing regressors. The equation of the four parameter sigmoid used to construct the timing regressor is given by:

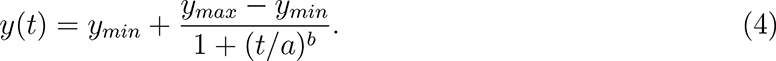

Here, y_min_ and y_max_ scale the vertical limits of the sigmoid, a controls the midpoint of the sigmoid, and b influences the slope of the sigmoid. The slope of the sigmoid at the midpoint is given by *−*b/4a. The parameters used in the full model were: y_min_=-1, y_max_=0, a = 4, b = *−*5.

The timing regressor for each session was mean-centered.

**Figure 10:**
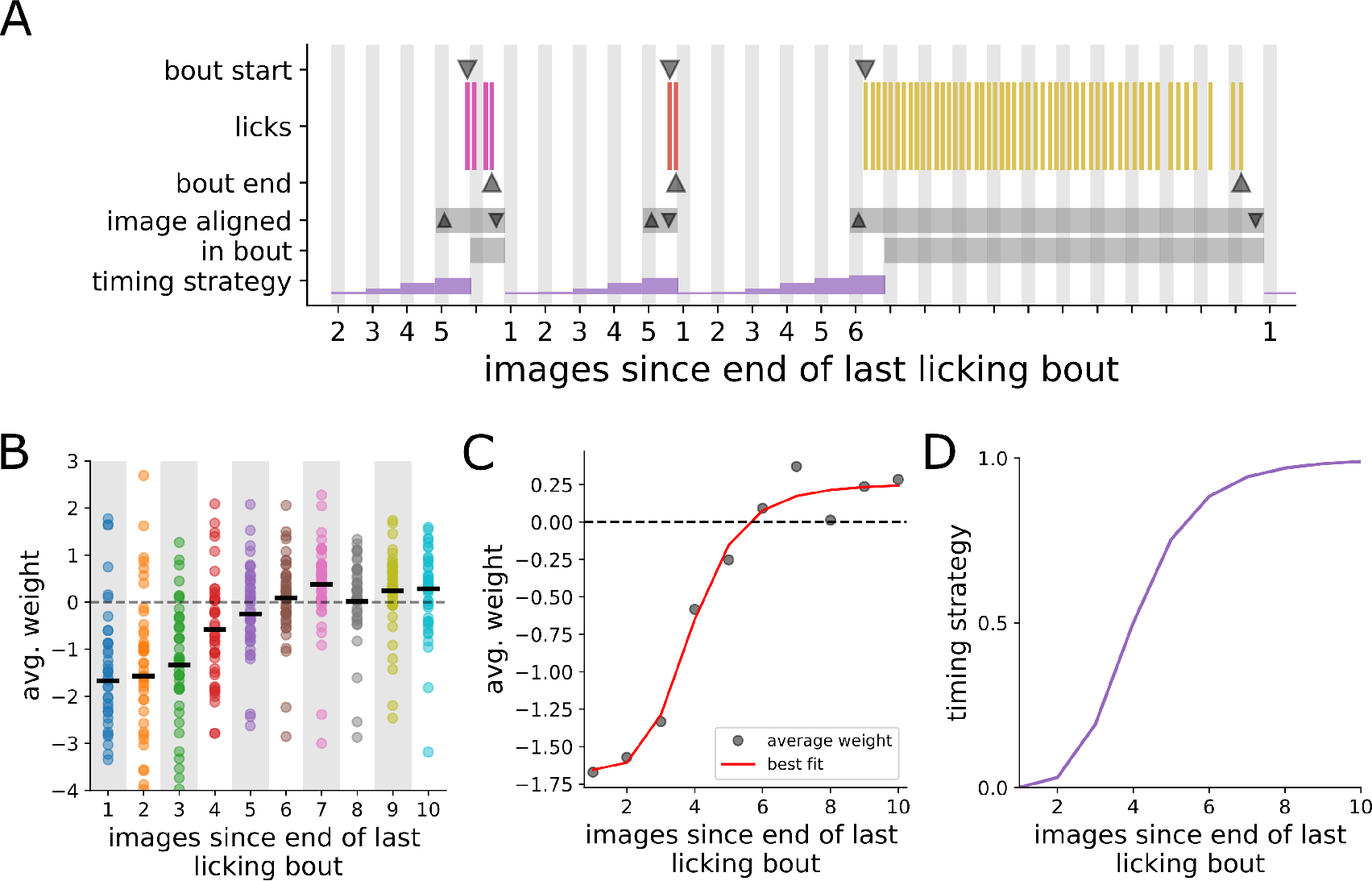
Constructing the timing regressor. (A) Schematic illustrating how timing is measured. Shaded bars indicate the time of stimulus presentations. Tick marks indicate the time of each lick. Individual licking bouts are colored separately. Down arrows (▽) indicate the start of each licking bout. Up arrows (*△*) indicate the end of each licking bout. Our model predicts whether the mouse will start a licking bout on each image presentation. Therefore images where the mouse was already in a licking bout are excluded from the fitting process. Consequently, our timing regressor starts measuring how many images have been presented since the end of the last licking bout starting at 1. The timing strategy is a sigmoidal function of time since the end of the last licking bout. Note the timing strategy is undefined on image when the mouse was already in a licking bout. (B) Average weight of each timing regressor (dots, n=45 sessions) Black bars indicate the average across sessions. (C) A four parameter sigmoid was fit to the average regressor weights from panel A (gray dots). The best fitting sigmoid is shown in red. (D) The timing strategy uses the midpoint and shape parameters from panel C, but scales the sigmoid to unit height. The strategy vector is later mean-centered.

## Supplemental Note 4 - Model validation

**Figure 11:**
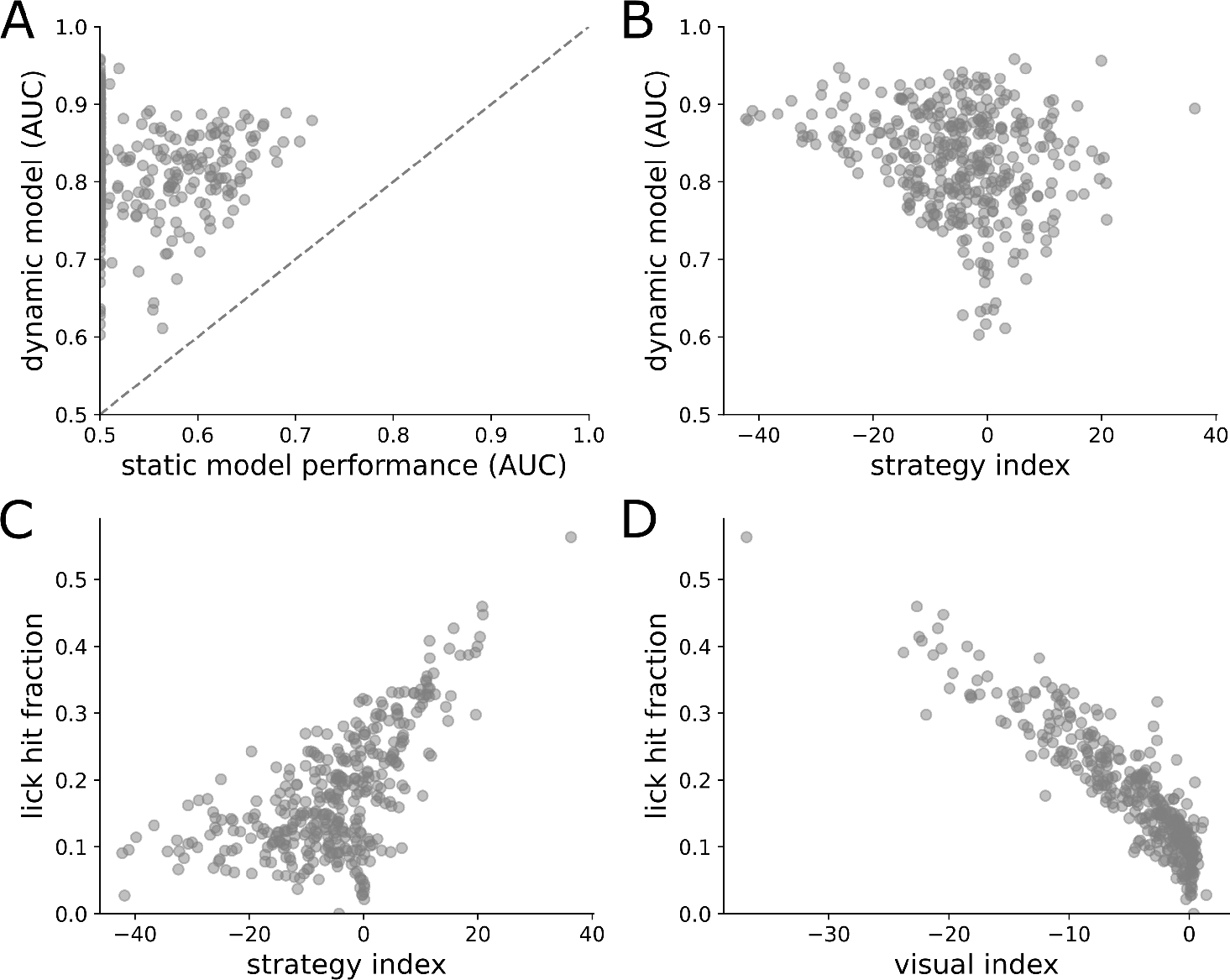
Model validation. (A) Scatter plot of area under ROC curves for each session for the dynamic model compared to static logistic regression (n=382 imaging sessions). Dashed line marks unity. (B) Scatter plot of area under ROC curves for each session for the dynamic model compared to the strategy index. (C-D) The lick hit fraction is the fraction of licking bouts that resulted in a reward. (C) Lick hit fraction compared to the strategy index. (D) Lick hit fraction compared to the visual strategy index.

## Supplemental Note 5 - Strategy characterization

**Figure 12:**
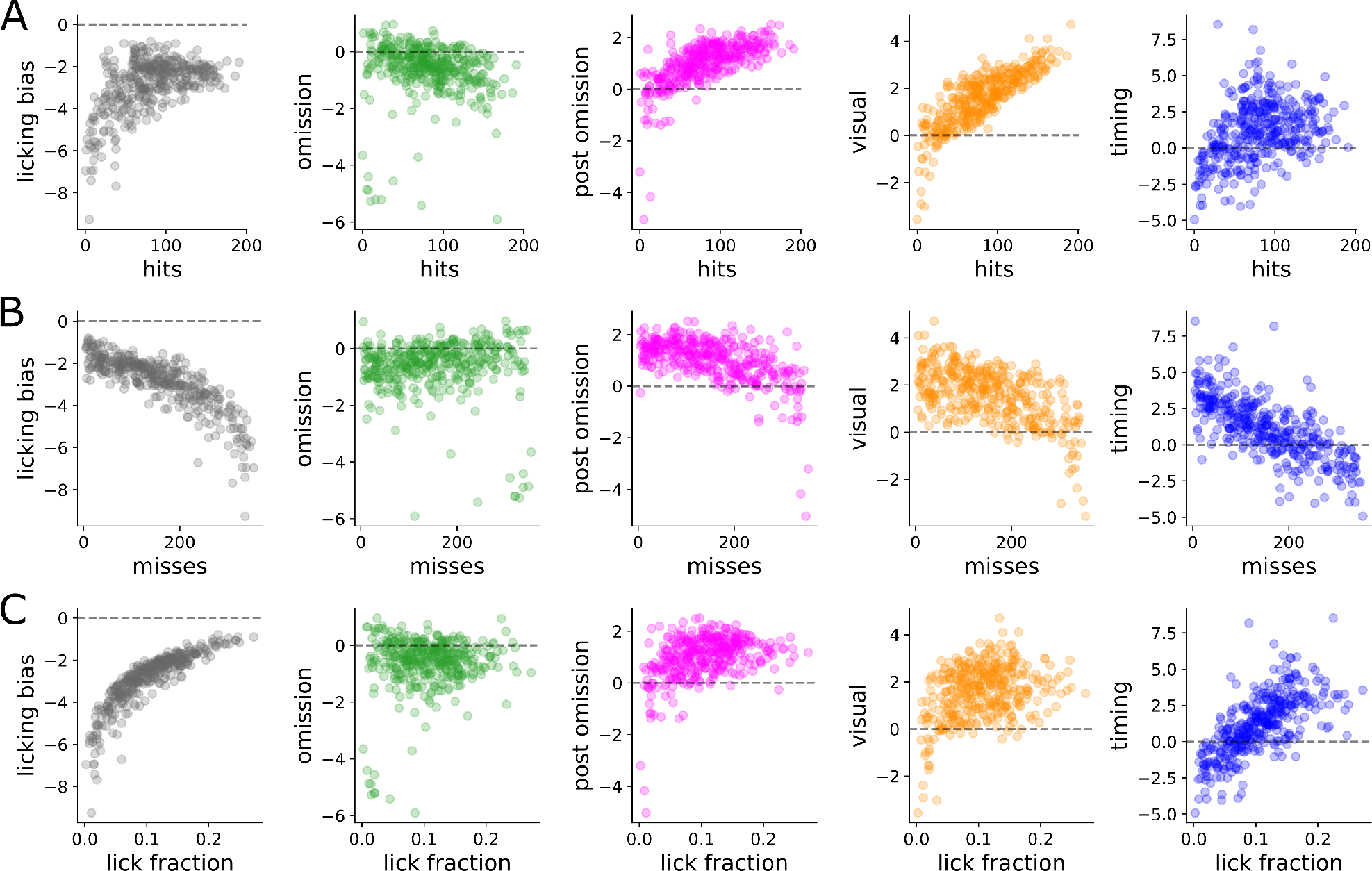
Average strategy weights are correlated with task events. Scatter plot between the average weight of each strategy and task events. Hits (A) are image changes with a reward, misses (B) are image changes without a reward, and lick fraction (C) is the fraction of images with a lick bout start.

## Supplemental Note 6 - Principal Components Analysis

To assess the variability of strategies across our behavioral dataset we performed PCA on the matrix of strategy indices containing 382 imaging sessions. Examining the component vectors we find that the first component is primarily aligned with the timing strategy, while the second is primarily aligned with the visual strategy. We find that these top two components contain 99.04% of the total variance (72.13% and 26.90%, respectively). Indeed, 98.24% of all variance in contained in just the timing and visual strategy indices. This finding motivates our use of the strategy index, which is simply the difference between the visual and timing indices. The strategy index contains 59.95% of the total variance, and has a strong correlation with the top principal component (R^2^ = 0.88).

**Figure 13:**
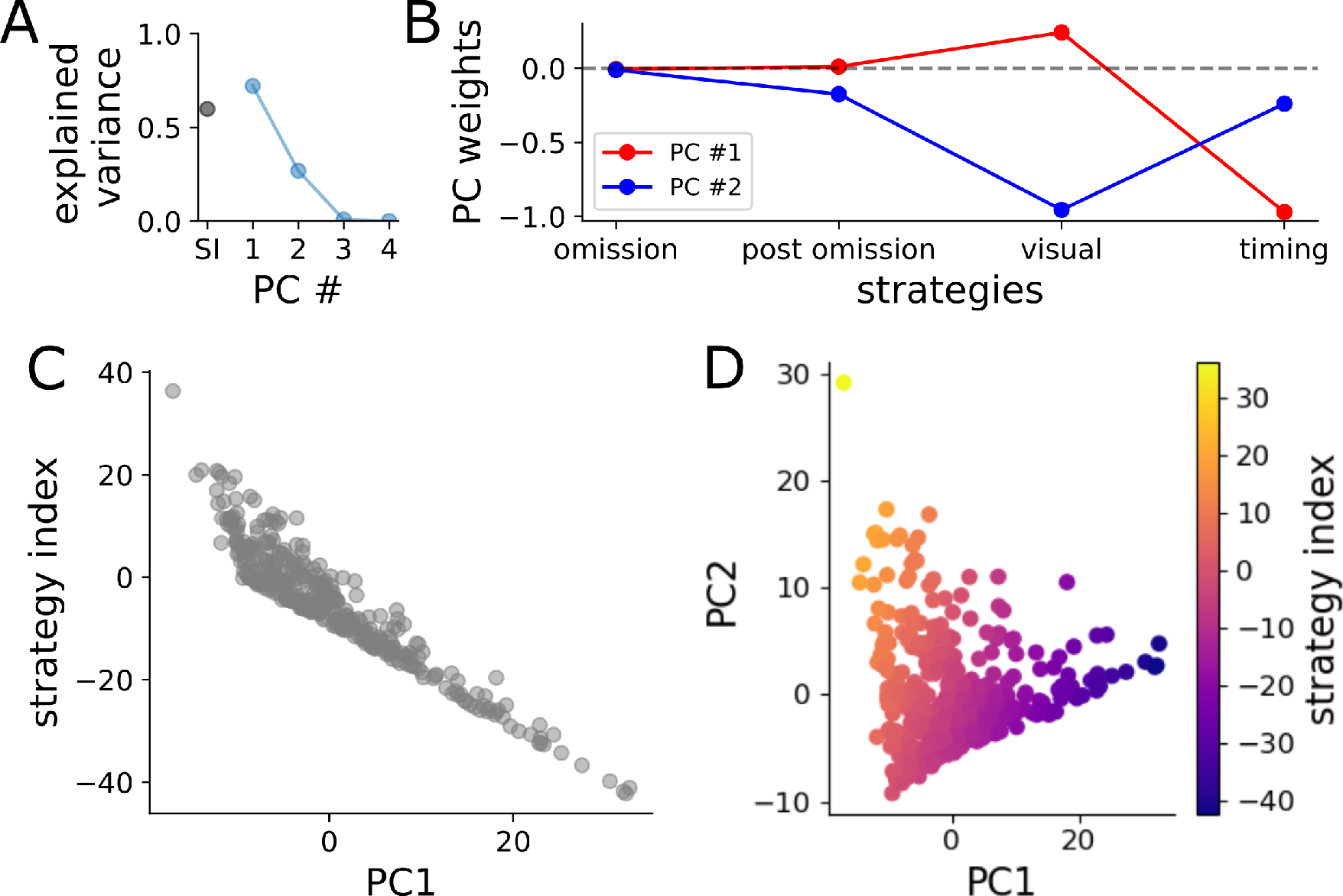
Principal Components Analysis (PCA) on strategy index. (A) Variance along each principal component (PC #), as well as the variance along the strategy index (SI). (B) The top two principal components are aligned with the timing and visual strategies, respectively. Scatter plot between each session projected onto the first principal component and the strategy index. (D) All sessions (n=382 imaging sessions) projected onto the first two principal components and colored by the strategy index.

## Supplemental Note 7 - Strategy over training

To examine how strategy preferences emerge with training, we classified mice by their dominant strategy (visual or timing) based on their behavior during imaging. We then fit our strategy model to the training sessions and examined how their strategy preferences changed over training. The training pipeline consisted of 7 stages before imaging:

- Training 0 - Mice learned to lick for rewards
- Training 1 - Mice earned rewards when a static grating changed orientation
- Training 2 - Static gratings were interleaved with a gray screen
- Training 3 - Static gratings are replaced with natural images
- Training 4 - Rewards decrease in size, and free rewards are no longer given when the mouse misses 10 changes in a row
- Training 5 - Performance must be consistently above a minimum threshold
- Habitation - Mice performed the task on the imaging rig
- Imaging - Mice performed the task with familiar images, then with novel images. Imaging was also performed during passive viewing of the same stimulus, which was not analyzed here.

We did not fit the model to training stages 0 and 1 because the stimulus was continuous and not periodically presented. In general we find strategy preferences emerge slowly over the training stages. Visual dominant mice significantly increased their use of the visual strategy over training (Fig. 13B), while decreasing their use of the timing strategy (Fig. 13C). Timing dominant mice showed less change in strategy use over training. Both strategies show decreases in the number of licking bouts (Fig. 13G), increases in the fraction of licking bouts that result in a reward (Fig. 13H), increases in the number of missed image changes (Fig. 13E), decreases in the fraction of the session they are engaged (Fig. 13F), while maintaining or slightly increasing the rewards per session (Fig. 13D). As mice increase their lick hit fraction, they need to lick less often to earn the same number of rewards. By increasing their lick hit fraction, this means they are licking on image changes more often rather than licking early which delays the next image change. Increasing their lick hit fraction means they can earn more rewards in less time, and thus they disengage earlier and miss more image changes while they are disengaged.

**Figure 14:**
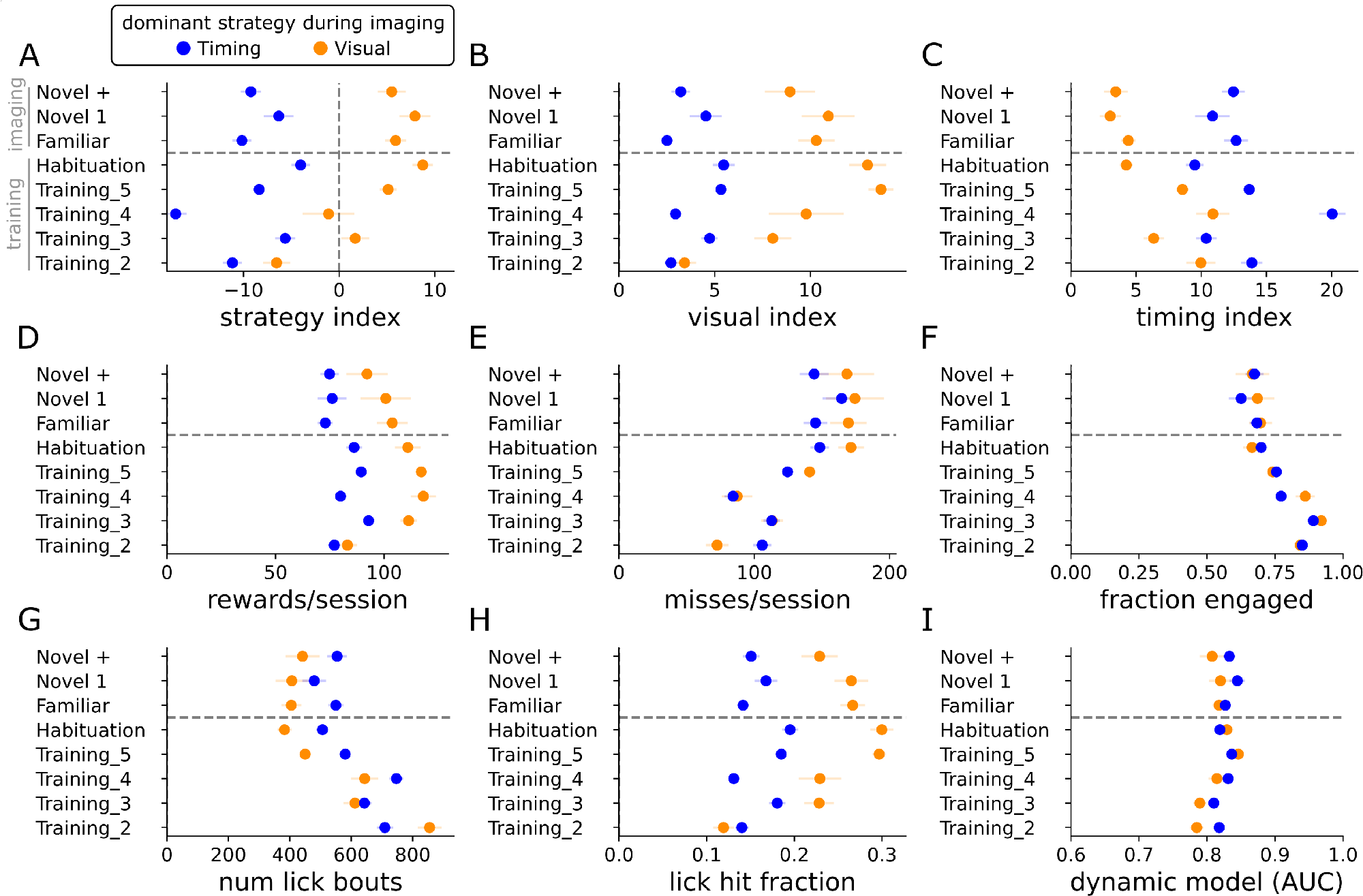
Strategy over training. Metrics as described in Supplemental Note 1. Each dot shows mean +/- SEM for each strategy group.

## Supplemental Note 8 - Strategy behavior over time

**Figure 15:**
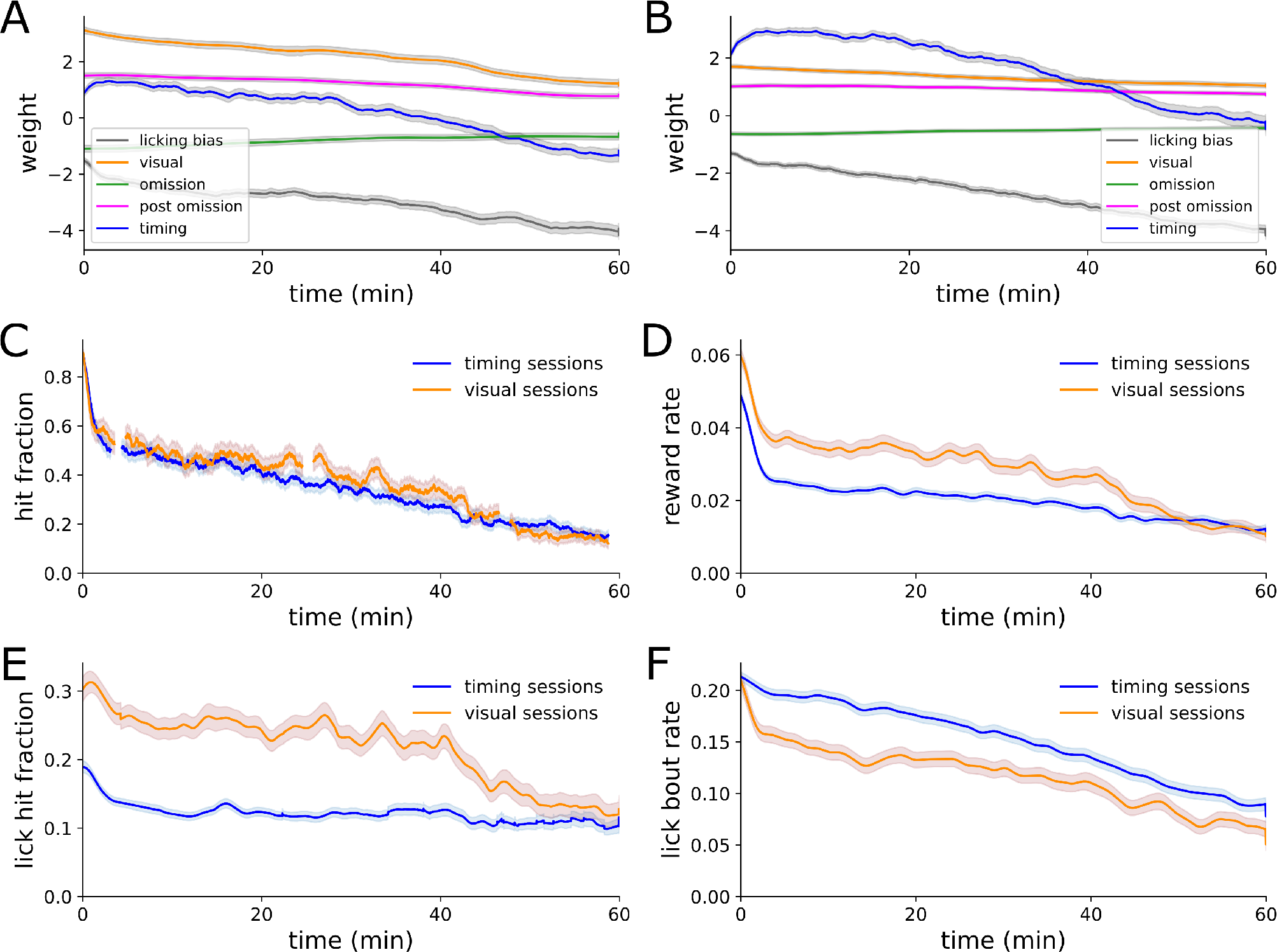
Strategy behavior over time. (A) Average strategy weights over time for visual strategy sessions (n = 116 sessions). (B) Same as A but restricted to timing strategy sessions (n=260 sessions). (C) Hit fraction split by visual or timing strategy sessions. (D) Reward rate split by visual or timing strategy sessions. (E) Lick hit fraction split by visual or timing strategy sessions. (F) Lick bout rate split by visual or timing strategy sessions.

## Supplemental Note 9 - Microcircuit dynamics

**Figure 16:**
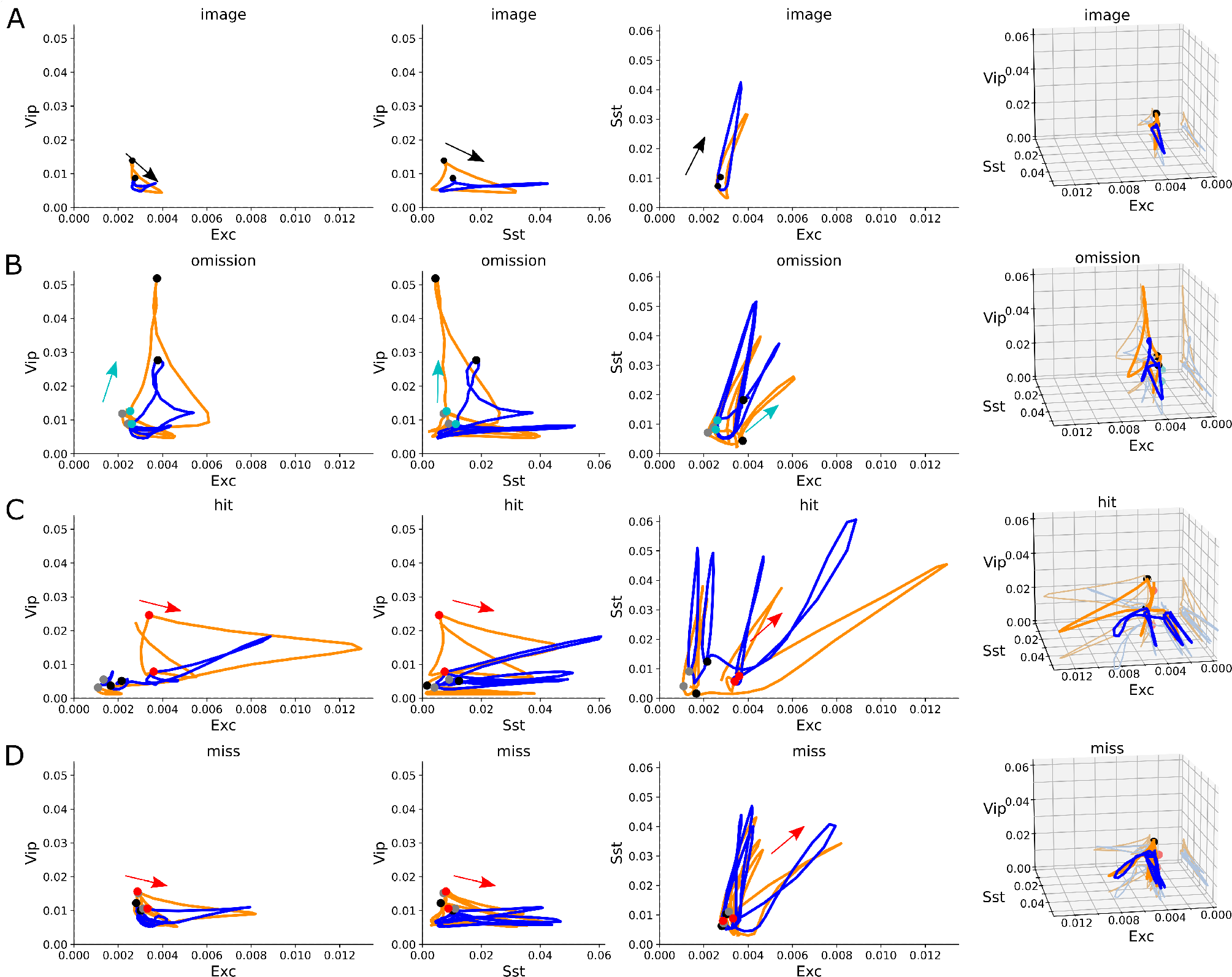
Microcircuit dynamics. Average population activity of different cell classes plotted against each other, similar to figure 5D. In response to (A) image repeats, (B) image omissions, (C) hits, and (D) misses. NOTE - need to polish, remove Vip-Sst vs Exc column, add 3D plots.

## Supplemental Note 10 - Running speed and task engagement

**Figure 17:**
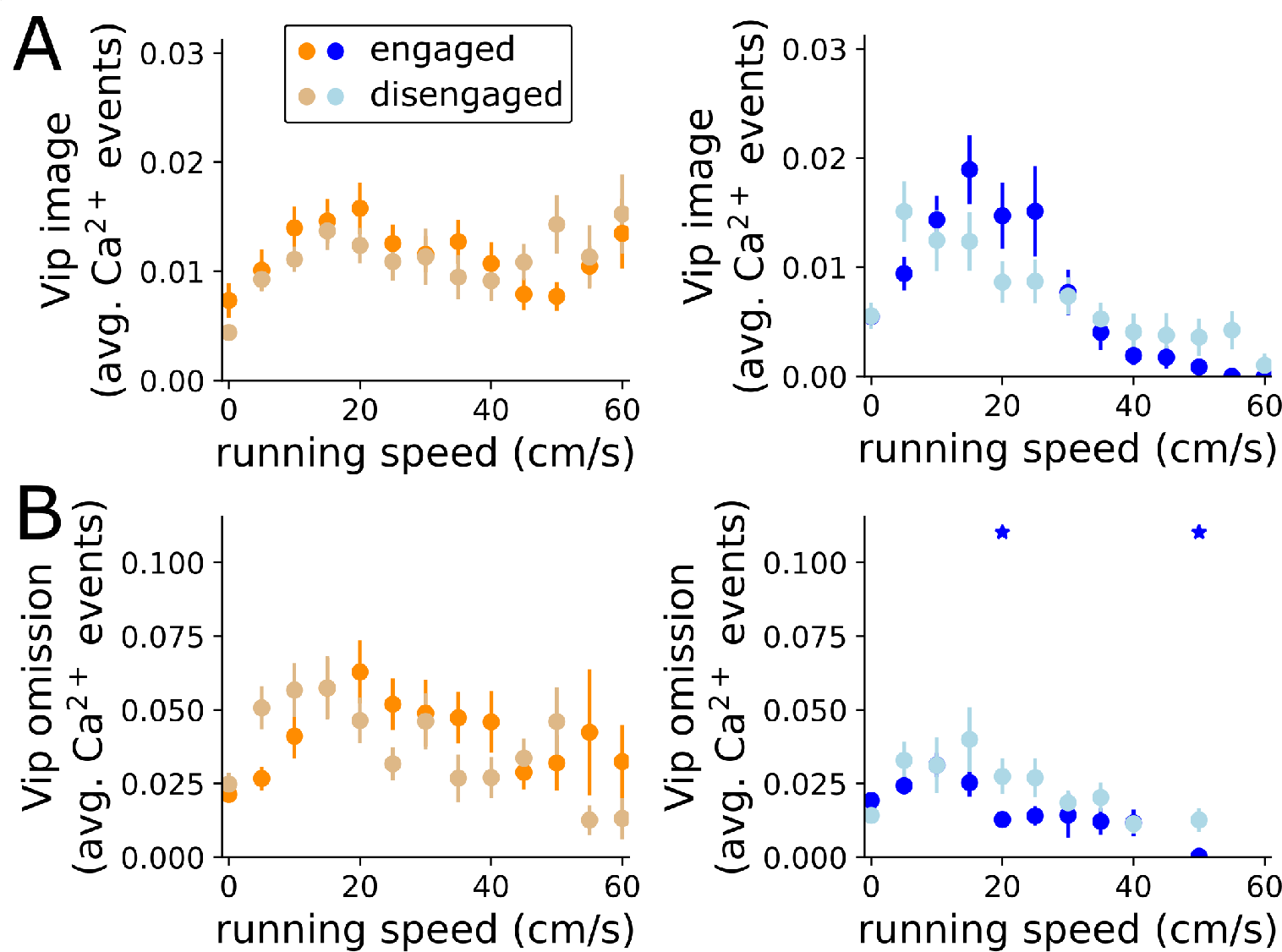
Running speed and task engagement. Similar to 4F,G. Vip activity in response to images and omissions across running speeds split by dominant strategy and task engagement. Stars indicates p <0.05 after a hierarchical bootstrap across imaging planes and cells, then corrected for multiple comparisons.

## Supplemental Note 11 - Transition to the novel image set

The mice were trained on one set of 8 images, we refer to this as as the familiar image set. After imaging during the familiar image set, the mice were transitioned to a new set of 8 images termed here as novel image set. We compare how this transition influenced mouse strategy. The familiar session is the last session using the familiar image set. Novel is the first exposure to the new image set. Novel+ is a repeated exposure to the new image set. The transition to novel stimuli is explored in-depth in Garrett et al. 2023. We make two notes here relevant to task strategy. First, there is a small but significant shift in the strategy index on novel sessions towards more visual strategy. Second, both strategies show the neural effects of novelty explored in Garrett et al. 2023. Thus any differences in strategy cannot explain the effects of novelty. Broadly, during the novel session we see increased Vip activation in response to image repeats, omissions, and before hits.

**Figure 18:**
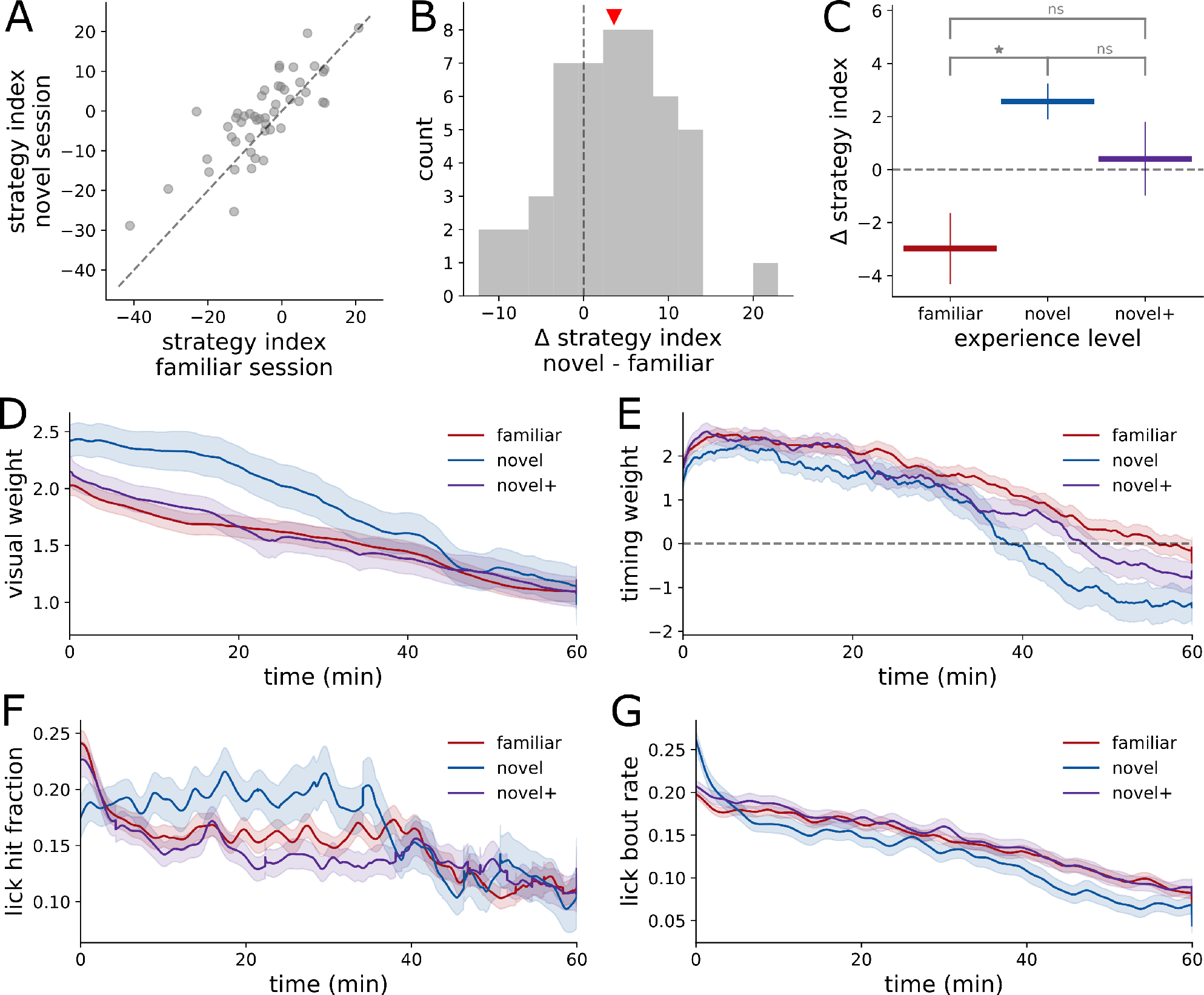
Stimulus novelty has a small influence on strategy. (A) Scatter plot of the strategy index for familiar and novel sessions. Each dot is a pair of sessions from the same mouse. (B) Histogram of the difference between strategy index on the novel session compared to the familiar session. (C) Average value of the strategy index across all mice relative to each mouse’s average strategy index value. Significance determined with a paired t-test, p<0.05. (D) Visual strategy weight over time split by experience level. (E) Same as D but for the timing strategy weight. (F) Lick hit fraction over time split by experience level. (G) Lick bout rate split by experience level.

**Figure 19:**
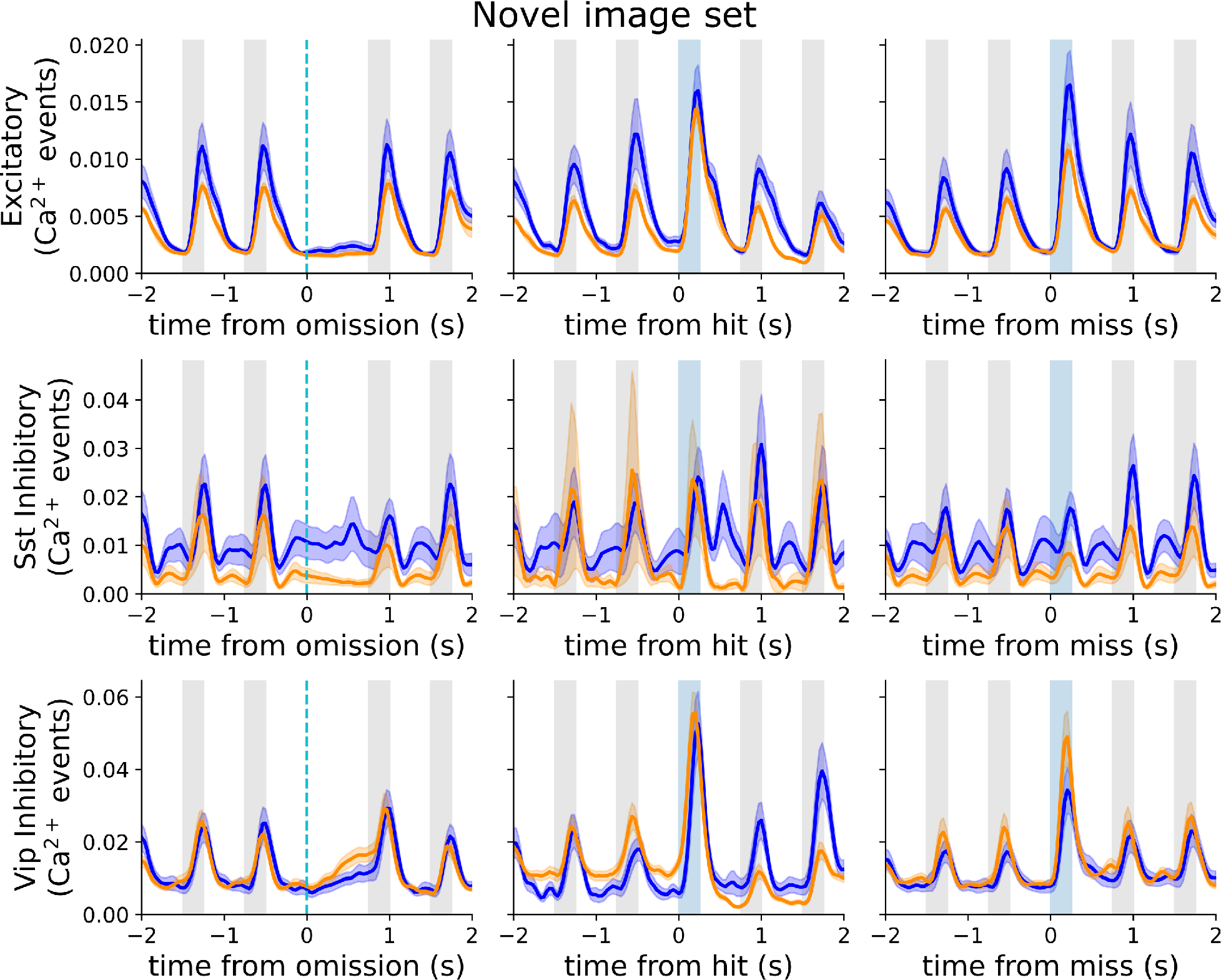
Both dominant strategies show robust changes to novel stimuli. Average population activity on sessions with novel stimuli for each cell class split by strategy aligned to either image omissions (left), hits (center), or misses (right). Compare with figure 4D, which show population activity on sessions with familiar stimuli.

